# Graph neural network modeling of receptor interaction kinetics from single-molecule imaging data

**DOI:** 10.64898/2026.07.08.737174

**Authors:** Khai Nguyen, Khuloud Jaqaman

**Affiliations:** Department of Biophysics, UT Southwestern Medical Center, Dallas, TX 75390, USA; Lyda Hill Department of Bioinformatics, UT Southwestern Medical Center, Dallas, TX 75390, USA

## Abstract

Single-molecule (SM) imaging (SMI)-based approaches have the powerful ability to capture receptor interactions – necessary for cell signaling – in their native live-cell environment. Yet, due to substoichiometric labeling, SMI generally provides only partial information on these interactions. We developed Deep-FISIK, which utilizes graph neural networks and multi-head attention for message-passing, to predict from SMI data the kinetics of homotypic interactions of the full receptor system. The input to Deep-FISIK are the SM detections in SMI experiments, without the need for explicit tracking. Thus, Deep-FISIK is compatible with labeling a higher fraction of receptors in the SMI experiments, increasing the prediction accuracy of the interaction kinetics parameters. Deep-FISIK’s performance is robust in the presence of a variety of deviations from the training data, indicating Deep-FISIK’s applicability to many receptor systems and SMI experiments.

## Introduction

Cell surface receptor interactions are critical for transmembrane signal transduction and cellular signaling in response to external stimuli^1–3^. Many receptors, including receptor tyrosine kinases^4–10^, G-protein coupled receptors^11,12^, immune cell receptors^13,14^, cytokine receptors^15,16^, among others^17,18^, undergo homotypic and heterotypic interactions that mediate and regulate their signaling function. Therefore, to achieve a quantitative understanding of cell signaling and its spatiotemporal regulation, it is important to quantify the kinetics (i.e., association and dissociation rate constants) of receptor interactions.

Quantifying the kinetics of molecular interactions, especially of cell surface receptors, in their native cellular environment is a challenging task. Traditional biophysical techniques, such as calorimetry^19^ or surface plasma resonance^20,21^, are performed *in vitro* and are not fully representative of the native cellular environment, especially of the lipid plasma membrane environment. These techniques might not be even applicable to cell surface receptors, which must reside in a relatively two-dimensional lipid environment. As such, the interaction kinetics obtained through traditional biophysical techniques (if possible at all) may not reflect the actual interaction kinetics of receptors on the cell surface^13,22–26^.

Microscopy offers routes to investigate molecular interactions in their native cellular environment^27,28^. One popular technique is dual-color fluorescence cross-correlation spectroscopy (FCCS)^29,30^. FCCS measures the concentrations of the species of interest, thus yielding the equilibrium dissociation constant K_d_ of their interactions. However, K_d_ is the ratio of the dissociation rate constant to the association rate constant, and the same K_d_ can result from fast kinetics or slow kinetics. Recent studies suggest that signaling can depend on the interactions’ dissociation rate^31–33^; thus, it is important to measure the interaction kinetics and not only K_d_. Additionally, many systems may be not at equilibrium^33,34^, in which case K_d_ is not applicable.

In order to calculate kinetics, temporal information on the interactions of interest must be obtained. Thus far, this has been done primarily for special scenarios. For example, fluorescence recovery after photobleaching (FRAP) can be used to estimate interaction kinetics when one of the interaction partners is immobile^35–37^. Ensemble FRET has been used to estimate the interaction kinetics between two proteins, however this required the micro-injection of one of the proteins into cells to create transients used for rate derivation^38^. Overall, determining receptor interaction kinetics, i.e., association and dissociation rates constants, in the cellular environment has remained a major challenge.

Other microscopy techniques that investigate molecular interactions in their native cellular environment and readily yield temporal information necessary to calculate interaction kinetics are those based on live-cell single-molecule imaging (SMI), including one-color, multi-color and SM FRET^9,17,34,39–45^. By labeling a subset of the molecule(s) of interest in live cells, such that individual molecules (or potentially a pair or triplet together) are distinguishable from each other, SMI techniques allow the capture of individual association and dissociation events between the imaged molecules. However, because of labeling only a subset of the molecule(s) of interest – necessary because of the resolution limit of the light microscope, which is ∼200 nm – many interaction events go undetected in SMI experiments, resulting in only a partial capture of the population interaction events.

Previous work in our lab demonstrated that combining SMI with mathematical modeling allows the inference of the full population interaction kinetics from the subset of interaction events observed via SMI^46^. The approach was termed “Framework for the Inference of in Situ Interaction Kinetics” (FISIK). However, FISIK had two main limitations: 1) the imaged molecules must be tracked, which may introduce errors that then propagate throughout the rest of the FISIK calculations; and 2) there is a conflict inherent to FISIK between the labeled fraction required to capture enough molecular interactions to infer their kinetics (higher is better) and the labeled fraction required for accurate tracking (lower is better).

To overcome these limitations, here we used graph neural networks (GNN) to predict molecular interaction kinetics directly from the graph of spatiotemporal connections between the detected molecules in SMI data, without the need for particle tracking. GNNs are powerful for this goal, as SMI data are readily depicted as a graph, where the graph nodes represent the detected molecules at each time point and the graph edges represent their connections in space and time. Deep learning methods, including GNNs, have been successfully applied recently to SMI data, but to predict only the diffusion properties of molecules, and not their interactions^47–50^ . To the best of our knowledge, our work – termed Deep-FISIK – is the first to employ deep learning to predict molecular interaction kinetics in situ from SMI data.

## Results

### Deep-FISIK predicts the maximum oligomeric state of the system and subsequently the SM system properties, including interaction kinetics

In order to predict the kinetics of molecular interactions, the types of interactions in a system of interest must be first known. In the case of homotypic interactions, this meant knowing the maximum oligomeric state in the system (Ν_M_), as this dictated the interaction parameters to be predicted. Note that Ν_M_ encompasses all of the oligomeric states that lead up to it. For example, Ν_M_ = 4 implies that a system contains monomers, dimers, trimers, and tetramers. Thus, starting with the graph of spatiotemporal connections between detected particles in an SM image series (graph nodes = simulated/detected particles, graph edges = potential connections between particles at different timepoints; Fig. 1A), we designed Deep-FISIK to consist of two steps:

**Figure 1.**
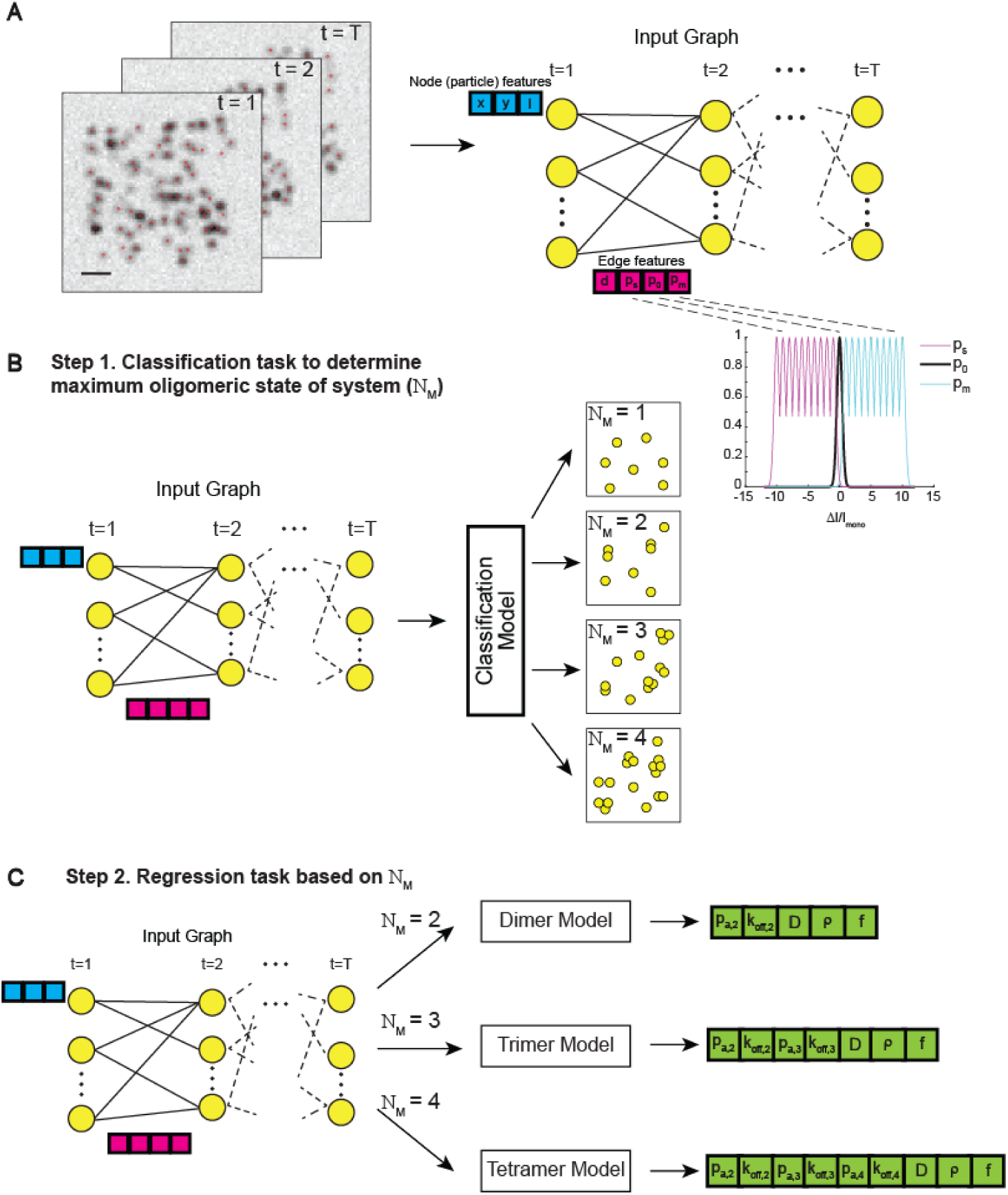
Deep-FISIK workflow to predict for a system of interacting molecules its maximum oligomeric state and interaction kinetics from SMI data. **(A)** Several time points of an SMI movie with detected SM particles overlayed in red (left). Image color is inverted for visual clarity. Scale bar, 1 µm. The detected particles over time are converted into a graph of spatiotemporal connections (right), used as input for the Deep-FISIK GNNs. Each detected particle is represented by a graph node and the connections between particles at different timepoints are represented by graph edges. Node features (x-coordinate (x), y-coordinate (y) and intensity (I) per detected particle) are in cyan. Edge features (distance between connected particles (d), and the probabilities of splitting (p_s_), no change in oligomeric state (p_0_), and merging (p_m_) when going from one particle to the connected particle) are in magenta. Inset shows p_s_, p_0_, and p_m_ as a function of the intensity difference between two connected nodes (ΔI) relative to the intensity of a monomer (I_mono_) (Eqs. 8-10). **(B)** Step 1 of Deep-FISIK, namely the classification task, to determine the maximum oligomeric state (Ν_M_) observed in a molecular system. For example, if a system contains monomers, dimers, trimers, and tetramers, then it is classified as a tetrameric system (Ν_M_ = 4). **(C)** Step 2 of Deep-FISIK, namely the regression task, to predict the various SM system properties, including interaction kinetics. The model used for the regression task depends on the system classification (Ν_M_) from Step 1.

Step 1. Classification task: Determine Ν_M_ (Fig. 1B). If Ν_M_ is known a priori, the classification task can be skipped.

Step 2. Regression task: Predict the SM system-level properties, based on the identified Ν_M_ (Fig. 1C). This includes the molecular interaction parameters, specifically the association probabilities *p*_a,ν_ (which reflect the propensity/readiness of molecules to interact when they are within each other’s proximity^46^) and dissociation rate constants *k*_off,ν_, both for ν = 2…Ν_M_, as well as general system properties, specifically molecule density *ρ*, labeled fraction *f*, and diffusion coefficient distribution mean *D* (averaged over all particles as a system-level property).

For both tasks, the architecture of the Deep-FISIK GNN models was comprised of encoder, message-passing, and decoder blocks (Fig. S1A), where multi-head attention (MHA) was used for message passing^51–53^ (Fig. S1B) (See Methods for more details). The models had two hyperparameters – the features dimension after embedding (encoder block) and the number of message-passing layers – which we optimized for each of the two tasks separately using hyperparameter tuning (discussed in the following two sections). Note that, because ground-truth interaction kinetics are generally not available for experimental SMI data, the GNN models were trained using simulated SMI datasets generated across a broad range of interaction parameters and other SM system properties, enabling supervised learning of the relationships between the graph structure and SM system properties, including molecular interaction kinetics.

In addition to the models’ architecture, it was important to employ node and edge features in the input graphs that were relevant for the properties to be predicted by Deep-FISIK. For the node features, we used the particles’ x- and y-coordinates and intensity. For the edge features, we used the Euclidean distance between connected particles, as well as intensity-based features that mapped the intensity difference between connected particles into a probability of oligomeric state change, thus reflecting molecular interactions (Fig. 1A inset). Briefly, similar intensities between two connected particles implied that no interaction occurred in going from one particle to the connected particle, an increase in intensity implied an association/merging event, and a decrease in intensity implied a dissociation/splitting event (see Methods for more details). These intensity-based features, which explicitly modeled changes in oligomeric state, were closely connected to the purpose of Deep-FISIK to predict interaction kinetics.

### Deep-FISIK identifies maximum oligomeric states of 1 to 4 with 99-74% true positive rate in pure simulated systems

The first step of Deep-FISIK was to determine the maximum oligomeric state of the system of interest, if not known a priori (Fig. 1B). To optimize the Deep-FISIK model architecture for this classification task, we trained 9 models with different combinations of number of node and edge features after embedding (48, 72 and 96) and number of message-passing layers L (6, 8, 11) (Fig. S2A). We did this initial training and testing on pure simulations data (5000 graphs per N_M_, N_M_ = 1-4), so as not to complicate the task with the limited resolution and noise of light microscopy images (to be done later). Following training, we validated the classification models of Deep-FISIK using unseen data, from which we calculated the confusion matrix between the systems with different Ν_M_.

To generate training and validation datasets for Deep-FISIK, we employed a stochastic model of molecular diffusion and interactions in 2D (Fig. 2A, B). Briefly, molecules (all of one type) with density *ρ* underwent free diffusion, where each molecule had a diffusion coefficient *D*_SM_ drawn from a normal distribution with specified mean *D* and standard deviation *σ*_D_ = 0.15*D*. Molecular interactions were controlled by association probabilities *p*_a,ν_ and dissociation rate constants *k*_off,ν_ , ν = 1 … N_M_. To mimic the substoichiometric labeling of SMI data, a fraction *f* of the molecules was labeled, leaving the remaining fraction (1- *f*) invisible^46^. See Methods for more details, and Table 1 for the ranges of parameter values employed for Deep-FISIK model training. The goal of the classification task was to predict N_M_, while the goal of the regression task (next section) was to predict *ρ*, *D*, *f*, *p*_a,ν_, and *k*_off,ν_ (ν = 1 … N_M_).

**Figure 2.**
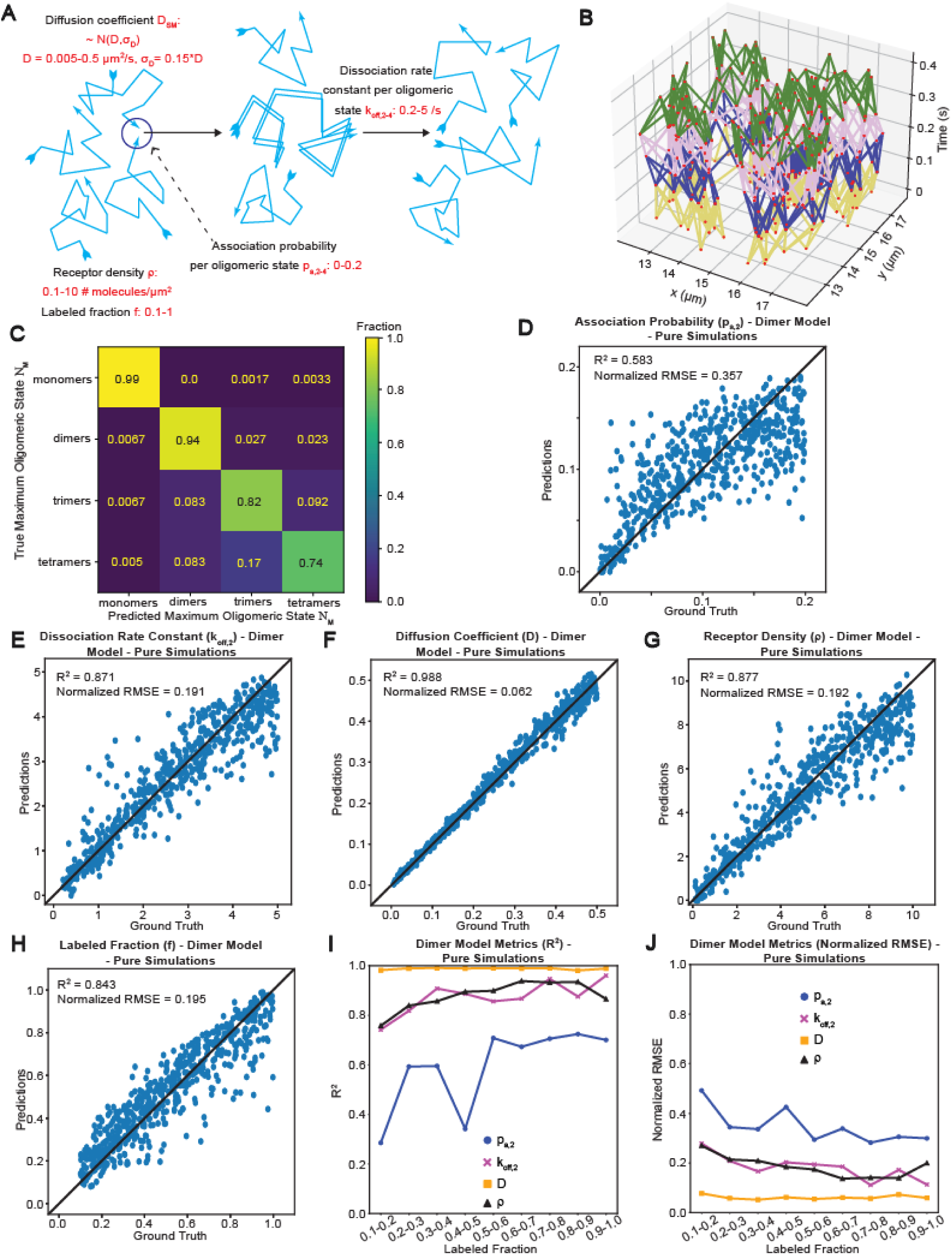
For pure simulated systems, Deep-FISIK predicts maximum oligomeric state with 74-99% true positive rate and dimeric SM system properties with relative error of 6-36%. **(A)** Illustration of the model used for simulating systems of interacting molecules. The schematic shows the critical SM system properties (model parameters), which Deep-FISIK then predicts, together with their ranges as used to generate training data for Deep-FISIK. **(B)** Visualization of the first 5 time points (0-0.4 s) of an input graph from one simulation. The simulated positions are shown as red dots. The edges connecting them are shown as colored lines going from yellow to blue to pink to green. **(C)** Confusion matrix displaying the performance of the pure simulations classification GNN model, as obtained by applying the classification model to unseen data (5000 and ∼600 graphs per maximum oligomeric state (i.e., 20,000 and ∼2,400 graphs in total) used for training and validation, respectively). **(D-H)** Scatterplots of predicted SM system properties vs. their ground truth values for the pure simulations Ν_M_ = 2 regression model, as obtained by applying the model to unseen data of dimeric systems. The five properties are the dimer association probability **(D)**, dimer dissociation rate constant **(E)**, diffusion coefficient distribution mean **(F)**, molecule density **(G)**, and labeled fraction **(H)**. Black straight line in each panel shows the unity line of perfect predictions. The pure simulations Ν_M_ = 2 regression model was trained on 30,000 graphs and evaluated on 600 graphs. **(I, J)** Plot of R^2^ **(I)** and normalized RMSE **(J)** displaying the performance of the pure simulations Ν_M_ = 2 regression model as a function of labeled fraction.

**Table 1.** SM system properties used in simulations.

| Property | Value range |
| --- | --- |
| $D$ | 0.005 – 0.50 $\mu\text{m}^2/\text{s}$ |
| $\rho$ | 0.1-10 # molecules/$\mu\text{m}^2$ |
| $f$ | 0.1-1 |
| $N_M$ | 1-4 |
| $p_{a,v}$, $v=2 \dots N_M$ | 0-0.2 |
| $k_{off,v}$, $v=2 \dots N_M$ | 0.2-5/s |

The model with 11 layers and 72 embedded features had the overall highest accuracy in predicting N_M_ among the tested models (Fig. S2A, B). The model was able to identify monomeric, dimeric, trimeric and tetrameric systems with, respectively, 99%, 94%, 82% and 74% true positive rate (Fig. 2C). The occasional misclassification occurred primarily with neighboring Ν_M_ (e.g., Ν_M_ = 3 was occasionally confused for Ν_M_ = 2 or Ν_M_ = 4). There was little confusion between the systems with interactions (Ν_M_ > 1) and systems without interactions (Ν_M_ = 1). All in all, our results indicate that Deep-FISIK is able to identify molecular systems with molecular interactions (vs. those without interactions) and to distinguish between Ν_M_ of 1-4.

### Deep-FISIK predicts SM system properties in a pure simulated dimeric system with a relative error of 7-36%

After determining Ν_M_, the second step of Deep-FISIK was to predict the SM system properties (Fig. 1C), with those related to molecular interactions of particular interest. Because the number of interaction parameters depended on Ν_M_, we devised and trained separate GNN models for the cases of Ν_M_ = 2, 3 and 4. As with the classification model above, we performed this initial training and testing on pure simulations data (image data to be handled later). Following training, we validated the various regression models using unseen data, through which we quantified the prediction accuracy for the general system properties (*D*, *ρ*, *f*) and for the interaction parameters (*p*_a,ν_, *k*_off,ν_, ν = 2…4).

First, we focused on the Ν_M_ = 2 model, as homotypic interactions forming receptor dimers are very common among receptors^7,10,39,54,55^. As with the classification model, we performed hyperparameter tuning to determine the optimal combination of number of layers and number of embedded features (using 30,000 graphs; see Table 1 for parameter value ranges). We found that, for the regression task, the combination of 8 layers and 96 embedded features had the highest overall accuracy (Fig. S2A, C). While all properties were predicted well by the model (Fig. 2D-H), there were some differences in predictability between the different properties. *D* was predicted excellently, with an R^2^ = 0.99 between the predictions and their ground truth values, and correspondingly a normalized RMSE (relative error) = 0.06. Next came *k*_off,2_, *ρ* and *f*, which were predicted very well, with an R^2^ ≈ 0.85 and a normalized RMSE = 0.19. The most difficult property to predict was *p*_a,2_, which nevertheless exhibited overall good correlation between its predictions and their ground truth values, with an R^2^ = 0.58 and a normalized RMSE = 0.36, indicating reliable predictability.

In the original FISIK, it was observed that the inference of interaction kinetics improved as *f* increased^46^. Is this the case also for Deep-FISIK? To investigate this, we plotted the prediction metrics for the N_M_ = 2 model as a function of *f* (Fig. 2I, J). Indeed, we found that the R^2^ and RMSE generally increased and decreased, respectively, as *f* increased. This suggests that the loss of prediction ability as *f* decreased was inherent to the information contained in the data, as the lower the labeling the more interactions are missed, making this information more and more difficult to recover. Interestingly, also as with the original FISIK^46^, the performance of Deep-FISIK with *f* > 0.2 was substantially better than its performance with *f* < 0.2. Given these similar performance trends, a clear advantage of Deep-FISIK compared to the original FISIK is that Deep-FISIK does not require particle tracking, thus allowing the labeling of a higher fraction of molecules for SMI, as needed for more accurate prediction of their interaction parameters.

### Deep-FISIK requires larger training datasets to predict interaction kinetics of higher order oligomers

Some cell surface receptors undergo higher-order oligomerization, forming trimers or tetramers, necessary for signaling^6,56–58^. Therefore, we also trained Ν_M_ = 3 and Ν_M_ = 4 regression models (using the same model architecture as for the Ν_M_ = 2 model). Initially we used 30,000 graphs for each model for training, as for the Ν_M_ = 2 model (see Table 1 for parameter value ranges). Testing these models on unseen data, we obtained similarly good prediction accuracy for *D*, *ρ* and *f* as with the Ν_M_ = 2 model (Fig. S3). The prediction accuracy of *k*_off,2,_ however, deteriorated, becoming similar to *p*_a,2_ (the accuracy of which remained similar to the Ν_M_ = 2 model) (Fig. S3). As for the additional interaction parameters describing higher order interactions (trimerization and tetramerization), their prediction accuracy deteriorated as the interaction order increased. In other words, the prediction accuracy of *p*_a,3_ and *k*_off,3_ was worse than that of *p*_a,2_ and *k*_off,2_ by the same model, and the prediction accuracy of *p*_a,4_ and *k*_off,4_ was worse than that of *p*_a,3_ and *k*_off,3_ by the same model (Fig. S3).

We reasoned that the worse prediction accuracy of the interaction parameters of higher order oligomers was because there were less instances of higher order interactions within each simulation (e.g., to form a trimer, first a dimer must be formed). To test whether this was the case, we trained another Ν_M_ = 3 model, this time using 60,000 graphs. Indeed, the prediction accuracy of all properties by this model was higher than that of the model trained on 30,000 graphs, including the prediction accuracy of the interaction parameters (Fig. S4). These results indicate that Deep-FISIK can predict the interaction parameters of higher order interactions, however the model training requires more data to achieve equivalent accuracy to the prediction of lower-order interaction parameters.

### Overestimating Ν_M_ leads to lower prediction errors by Deep-FISIK than underestimating it

In the Deep-FISIK framework, the regression model used for predicting the interaction parameters depends on the results of the classification task (Fig. 1; assuming that Ν_M_ is not known a priori). If Ν_M_ gets misclassified, then an incorrect regression model will be utilized. What are the consequences of using an incorrect model for the regression task? To mimic this situation and investigate the consequences of misclassifying a system, we applied the Ν_M_ = 2 model (Fig. 2) to systems containing up to trimers (Fig. S5; emulating a case of underestimating Ν_M_) and the Ν_M_ = 3 model (the model trained on 60,000 graphs; Fig. S4) to systems containing only up to dimers (Fig. S6; emulating a case of overestimating Ν_M_). This testing was motivated by our observation that, when the classification model made an error, it mostly misclassified Ν_M_ by ±1 (Fig. 2C).

These tests revealed that applying the Ν_M_ = 3 model to a dimeric system (Fig. S6) has fewer negative consequences for property prediction than applying the Ν_M_ = 2 model to a trimeric system (Fig. S5). When applying the Ν_M_ = 2 model to a trimeric system, i.e., underestimating Ν_M_, the prediction trends suggest that the model sensed that there were more interactions than expected for a dimeric system, but as the model was restricted to dimers, its attempt to represent the more-than-expected interactions led to systematic biases in the property predictions. This manifested itself in overestimating *p*_a,2_ and underestimating *k*_off,2_ (Fig. S5A, B). It also manifested itself in overestimating *f*, as with higher *f* more interactions of a system would be visible, and simultaneously underestimating *ρ* (*f* and *ρ* tend to inversely correlate as the product of the two equals the observed density of receptors^46^).

In contrast, applying the Ν_M_ = 3 model to a dimeric system, i.e., overestimating Ν_M_, was less consequential for prediction accuracy, as a dimeric system is in principle a subset of a trimeric system, specifically with *p*_a,3_ = 0. In this case, the predicted properties showed minimal systematic bias and correlated reasonably well with their ground truth values (but worse than when using the correct Ν_M_ = 2 model; Fig. 2D-H vs. Fig. S6). As for the artifactual trimer interaction parameters that got predicted in this case for the dimeric system, *p*_a,3_ was mostly predicted to be close to zero, as it should be (Fig. S6A, right). *k*_off,3_ is fundamentally undeterminable for a dimeric system, and interestingly its predicted values were concentrated in the above average part of the *k*_off_ range (Fig. S6B, right). This implies that the trimers “imagined” by the Ν_M_ = 3 GNN model were short-lived, which is as close as the Ν_M_ = 3 model could get to a dimeric system. All in all, these results suggest that, when Ν_M_ is uncertain, utilizing a higher N_M_ model yields overall more accurate results than utilizing a lower N_M_ model. A potentially advantageous strategy is to utilize multiple models, namely N_M_ = 2, N_M_ = 3, etc., and seek convergence for lower order interaction parameters and simultaneously an association probability close to zero for higher order oligomers.

### Deep-FISIK identifies maximum oligomeric states of 1 to 4 with 95-63% true positive rate in synthetic images

The above training and testing were performed on graphs from pure simulations, in which case every molecule position was known without errors; pure simulations contain no noise that would increase the uncertainty of molecule detection and localization, and they have no resolution limit that would prevent us from distinguishing between molecules no matter how close they are to each other. To mimic experimental data more closely and investigate Deep-FISIK’s performance under more realistic conditions, we generated image series from the above-described simulations, replacing each molecule position with a point-spread-function (PSF; approximated as a Gaussian with standard deviation = 122 nm) and adding background noise, such that the image signal-to-noise ratio (SNR, defined as (SM intensity – average background intensity) / (background intensity standard deviation) was in the range 4-7, similar to typical SMI data^9,59^ (Fig. 3A, Movies S1-S8, and Methods). We then detected and localized the particles in the images using u-track^60^ (Fig. 3A, Movies S1-S8), and used the detected particles to construct graphs for model training and evaluation (equivalent to what was done above). Because the properties of the detections from images were expected to be different from those of pure simulated positions, we trained a new classification model using the graphs generated from detections (using the same hyperparameters optimized for the pure simulations classification model).

**Figure 3.**
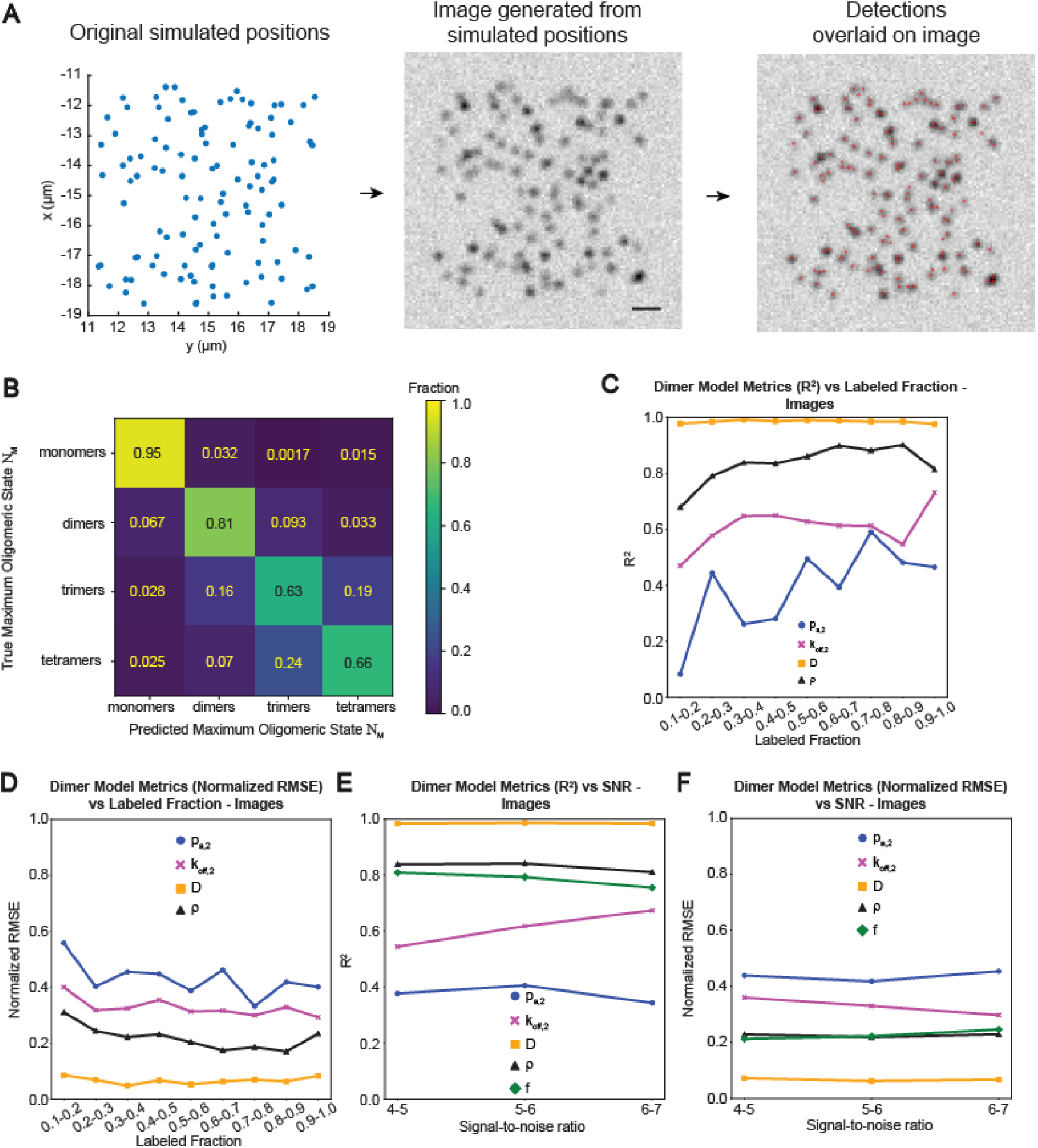
For images, Deep-FISIK predicts maximum oligomeric state with 63-95% true positive rate and dimeric SM system properties with relative error of 8-45%. **(A)** Starting with simulated positions (left), synthetic images are generated by replacing each simulated particle with a point spread function and adding background noise (center), after which particles are detected in the synthetic images using Gaussian mixture-model fitting (right). Scale bar, 1 μm. **(B)** Confusion matrix displaying the performance of the images classification GNN model, as obtained by applying the classification model to unseen image data (∼5,000 and ∼600 graphs per maximum oligomeric state (i.e., ∼20,000 and ∼2,400 graphs in total) used for training and validation, respectively, all generated synthetically). **(C, D)** Plot of R^2^ **(C)** and normalized RMSE **(D)** displaying the performance of the images Ν_M_ = 2 regression model for the various predicted SM system properties as a function of labeled fraction, as obtained by applying the model to unseen image data of dimeric systems. The model was trained on ∼30,000 graphs and evaluated on ∼600 graphs (all generated synthetically). **(E, F)** Plot of R^2^ **(E)** and normalized RMSE **(F)** displaying the performance of the images Ν_M_ = 2 regression model for the various predicted SM system properties as a function of image signal-to-noise ratio (SNR). Same model and data as in C, D.

The classification model for synthetic images (trained on ∼5000 graphs per N_M_, N_M_ = 1-4) trained well and was able to perform its task, as evaluated on unseen image data, albeit with slightly lower performance than the model trained on and applied to pure simulations (Fig. 3B, Fig. S7). The overall true positive rate of the classification model predictions was 77% (Fig. 3B; as compared to an overall true positive rate of 87% for the pure simulations). Like the pure simulations classification model, the images classification model distinguished well between systems of no interactions and systems with interactions (N_M_ = 1 vs. N_M_ > 1), and the cases of misclassifying N_M_ were largely confined to neighboring N_M_ (i.e., ±1).

### Deep-FISIK predicts SM system properties in a synthetic image dimeric system with a relative error of 8-45%

As with the pure simulations case, after the classification step Deep-FISIK applied to image data would perform a regression step to determine the SM system properties. Therefore, we also created an N_M_ = 2 regression model trained on synthetic images (using ∼30,000 graphs), generated from the above-described simulations using a similar process as the classification model (see Table 1 for parameter value ranges). We limited the images regression model to the N_M_ = 2 case as it was simpler and required less training data than the models containing higher order oligomers. As with the pure simulations regression model, the performance of the images regression model varied by SM system property. The prediction accuracy for *D*, *ρ*, and *f* was very similar to that of the pure simulations regression model (Fig. S7C-E vs. Fig. 2F-H). The prediction accuracy for *p*_a,2_ and *k*_off,2_, on the other hand, was worse than that of the pure simulations regression model. The R^2^ correlating the predicted values to their ground truth values was 0.61 and 0.38 for *k*_off,2_ and *p*_a,2_, respectively, as compared to 0.87 and 0.58 in the pure simulations case (Fig. S7A, B). Nevertheless, as with the pure simulations model, the prediction performance for all properties generally increased as the labeled fraction increased (Fig. 3C, D). It was very similar for the different SNRs (Fig. 3E, F).

It is not surprising that the two properties with worse performance in the synthetic images case are those related to molecular interactions; the detection and characterization of molecular interactions is compromised much more than other molecular properties, such as movement, by the limited resolution of light microscopy. The stable performance of the model across different SNRs supports the notion that the lower ability to predict *p*_a,2_ and *k*_off,2_ from images stems primarily from the limited resolution of light microscopy, and not from detection or localization errors due to limited signal (at least in the SNR range of 4-7). Of note, creating an images regression model allowed us to compare our model’s performance to a previously published GNN model that was focused on predicting molecular diffusion properties from SMI detections^49^. In this previous work, the model’s accuracy in predicting the diffusion coefficient (over a range of values that overlaps with the range employed in our study) was quantified using the mean absolute error (MAE), and was found to be 0.06^49^. In comparison, the MAE quantifying Deep-FISIK’s accuracy in predicting *D* was 0.01. This demonstrates that Deep-FISIK’s performance is comparable if not exceeding that of previous models in the case of predicting more accessible SM system properties, such as the diffusion coefficient, and highlights the challenge of predicting properties related to molecular interactions.

Interestingly, our results indicate that the predictions from images improve with increased *f*, at least up to *f* ≈ 0.7-0.8 (Fig. 3C, D). This sustained trend in images is somewhat counterintuitive, as labeling at higher fractions is expected to exacerbate the effects of the resolution limit. However, it appears that the additional information provided by higher *f* outweighs this limitation for *f* < 0.7- 0.8. These findings again highlight the strength of Deep-FISIK in that it does not require explicit particle tracking, thus allowing the higher labeling of molecules for SMI experiments, as needed for more accurate prediction of their interaction kinetics.

### Interaction kinetics parameter prediction is robust to variations in molecular diffusion coefficient distribution

Our models were trained on systems where the SM diffusion coefficient followed a normal distribution, *D*_SM_ ∼ N(*D*, *σ*_D_), where *D* was in the range 0.005-0.5 µm^2^/s and *σ*_D_ = 0.15*D*. However, the SM diffusion coefficient distribution observed in experimental data is often more complex^9,17,59,61,62^. Therefore, we tested the robustness of our trained models against *D*_SM_ distributions that deviated from this idealized distribution. We tested four scenarios: (i) *D*_SM_ follows a wider normal distribution, with *σ*_D_ = 0.25*D*; (ii) *D*_SM_ follows a log normal distribution, which, unlike the normal distribution, exhibits a long right tail, which is closer to what is often observed experimentally^9,17,59,62^; (iii) all molecules have the same *D*_SM_ = *D*; and (iv) *D*_SM_ follows an experimentally-derived distribution, specifically that observed for the cell surface receptor VEGFR2^59^. Using these different *D*_SM_ distributions, we simulated molecules diffusing and interacting to form (reversible) dimers (using the same model parameter ranges in Table 1). From these simulations we then generated images, detected the particles in the images, and constructed graphs to which we applied the images classification and regression models.

In all scenarios except for the experimental VEGFR2 distribution, model performance for both the classification and regression tasks was very similar to the baseline model performance (i.e., model performance when applied to data simulated using the *D*_SM_ distribution employed in model training) (Fig. 4A, B). In the experimental VEGFR2 *D*_SM_ distribution scenario, the classification true positive rate was somewhat lower, at 65% (Fig. 4A), as was the accuracy of *D* prediction (normalized RMSE of ∼0.3 vs. 0.07; Fig. 4B). Importantly, however, the prediction accuracy of the other SM system properties, including the interaction parameters, was comparable to the baseline. There was some reduction in prediction ability for *k*_off,2_ (normalized RMSE increased from ∼0.3 to ∼0.4), but the prediction ability of *p*_a,2_ was not affected (normalized RMSE remained ∼0.45). This uncoupling between *D* prediction and interaction kinetics predictions most likely stems from the fact that *D* is most heavily dependent on the distance edge feature, while the interaction parameters are primarily dependent on the intensity-based features.

**Figure 4.**
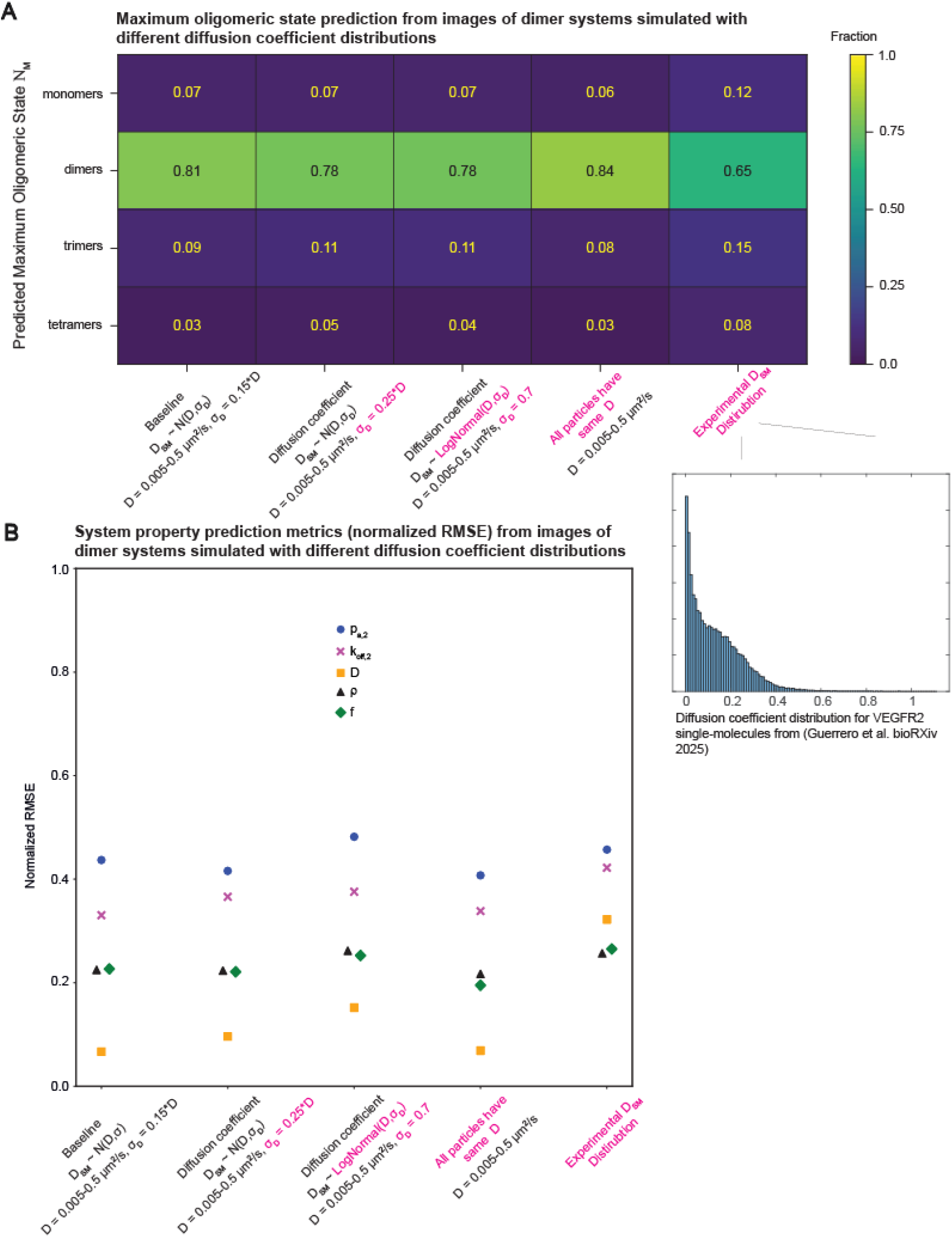
Deep-FISIK is largely robust when applied to systems with diffusion coefficient distributions differing from the training dataset. **(A)** Classification accuracy when applying the images classification model (same model as in Fig. 3) to image datasets of dimeric systems generated using various diffusion coefficient distributions: normal distribution but with different standard deviation, log-normal distribution, same diffusion coefficient for all particles, and SM diffusion coefficient sampled from experimental distribution from SMI of VEGFR2^37^. All systems are simulated to be dimeric, but the classification model can predict from monomeric to tetrameric. Dataset size = ∼600 graphs per diffusion coefficient distribution type. **(B)** Normalized RMSE plot for the SM system properties predicted by the images Ν_M_ = 2 regression model (same as in Fig. 3) for image datasets of dimeric systems generated with different diffusion coefficient distributions (same dataset as in A).

These results indicate that the Deep-FISIK GNN models are reasonably robust in the presence of deviations from the *D*_SM_ distribution employed for model training. If the *D*_SM_ distribution of a particular dataset deviates heavily from the training *D*_SM_ distribution (as seems to be the case for the VEGFR2 distribution tested here), then the *D* prediction might be inaccurate. In this case, it is advisable to retrain the Deep-FISIK models for that specific scenario, in order to improve the *D* prediction, and potentially that of other properties, such as *k*_off_. Note that our data-generation model assumed freely diffusing molecules – with *D*_SM_ following a normal or other distribution – but for some SM systems other diffusion types might be relevant as well^7,17,61^. The diffusion types relevant for a particular dataset, and/or the empirical *D*_SM_ distribution, can be obtained by SM particle tracking and diffusion analysis^47,63^, and thus can be readily compared to the predictions (and assumptions) of Deep-FISIK. Although the prediction of *D* is largely uncoupled from the predictions of the other SM system properties, agreement between the *D* predicted by Deep-FISIK and *D* obtained via other analysis methods (such as particle tracking) would lend confidence in the predictions of Deep-FISIK.

### Classification task of Deep-FISIK is largely robust to out-of-range SM system properties

For a comprehensive characterization of Deep-FISIK, we tested next Deep-FISIK’s performance when the SM system property values are outside of the training range. In general, neural networks cannot predict outside of their training range, i.e., they cannot extrapolate^64^, and thus it was important to investigate Deep-FISIK’s behavior in this regard. For these tests, we generated dimeric system image series (as above), where one or the other system property was out of range: (i) *p*_a_^>^: association probability in the range 0.2-0.3 (above the training range); (ii) *k*_off_^<^: dissociation rate constant *k*_off_ in the range 0.1-0.2/s (below the training range); (iii) *k*_off_^>^: dissociation rate constant in the range 5-10/s (above the training range); and (iv) *ρ*^>^: receptor density in the range 10-20 mol/μm^2^ (above the training range). In each case, the other properties had values within the training range, i.e., we tested the effect of property deviations one at a time.

To our surprise, the classification task of Deep-FISIK was quite robust in the presence of out-of-range SM system properties (Fig. S8A). The tested systems were correctly identified as dimers with a true positive rate similar to, or even better than, the baseline in the *p*_a_^>^, *k*_off_^>^ and *ρ*^>^ scenarios. Only in the *k*_off_^<^ scenario was the true positive rate lower (0.69 instead of 0.81), and interestingly in an explainable direction. Specifically, more systems were misclassified as trimeric in this case, which probably stemmed from the reduced number of dissociation events observed by the model when *k*_off_ was below the training range. Note that in the other scenarios, although the true positive rate was similar to the baseline, the misclassification also shifted in a manner consistent with the shifts in the SM system properties. Specifically, in the *p*_a_^>^ scenario (more association events) most misclassifications were to Ν_M_ = 3, and in the *k*_off_^>^ scenario (more dissociation events) most misclassifications were to Ν_M_ = 1. The higher true positive rate of the *ρ*^>^scenario was probably due to the increased occurrence of interaction events (both association and dissociation) when there were more molecules, i.e., the model had more data based on which it could predict.

In contrast to the classification task, the regression task of Deep-FISIK was not able to extrapolate, as expected. Out-of-range system properties were not predicted accurately (Fig. S7A, B, D, Fig. S8B). In addition, one property being out-of-range often affected the prediction accuracy of other properties. In particular, *p*_a,2_ and *k*_off,2_ were coupled, such that one being out of range negatively impacted the prediction of the other. Interestingly, *k*_off,2_ being out of range had a much higher impact on the prediction accuracy of *p*_a,2_ than the other way around. All properties were also badly predicted in the *ρ*^>^ scenario (in contrast to the classification task, which improved in this scenario).

All in all, these results indicate that the classification task is largely robust to out-of-range SM system properties, but the regression task is not (the latter as expected). Our training dataset covered a wide range of SM system property values, informed by previous studies^9,59,65,66^, in an attempt to circumvent this problem. Nevertheless, for a sanity-check of Deep-FISIK predictions, a prudent strategy is to combine Deep-FISIK with system perturbations and/or biological controls with known effects on the interaction kinetics (and other system properties) and to verify that Deep-FISIK’s predictions follow the expected changes.

### Deep-FISIK is robust to changes in imaging conditions as long as they are compatible with the SM system properties

Finally, we tested the robustness of Deep-FISIK against variations in imaging conditions. The classification and regression models were trained on simulations that were 15 s long and sampled at 10 Hz. However, there could be other biological systems or image acquisition setups with different imaging conditions. For example, if diffusion is faster or dissociation happens at a faster scale, then faster sampling would be required. Or, if dissociation happens at a slower scale, then slower sampling and longer total acquisition time would be required. Thus, we tested how robust Deep-FISIK is when the imaging conditions vary, as that affects the applicability of Deep-FISIK to a variety of biological systems and imaging setups. We tested Deep-FISIK for robustness against longer or shorter acquisition times, and faster or slower sampling rates.

In terms of changes in total acquisition time, we found that increasing the total acquisition time had largely no negative effect on Deep-FISIK’s performance, for both the classification task and the regression task, except for a slight worsening of the prediction accuracy for *k*_off,2_ (Fig. 5A, B). Decreasing the total acquisition time, however, negatively impacted the classification task, where classification accuracy dropped from 0.81 to 0.68 (Fig. 5A). Similar to increasing the total acquisition time, decreasing the total acquisition time slightly worsened the prediction accuracy of *k*_off,2_, while the other SM system properties were not impacted (Fig. 5B). All in all, these results suggest that the regression task is reasonably robust to changes in total acquisition time, while the classification task is negatively impacted when the total acquisition time is decreased. Interestingly, more systems got misclassified as trimeric instead of dimeric in the case of shorter total acquisition time (Fig. 5A), potentially because a shorter acquisition time led to fewer dissociation events in the system observation time window.

**Figure 5.**
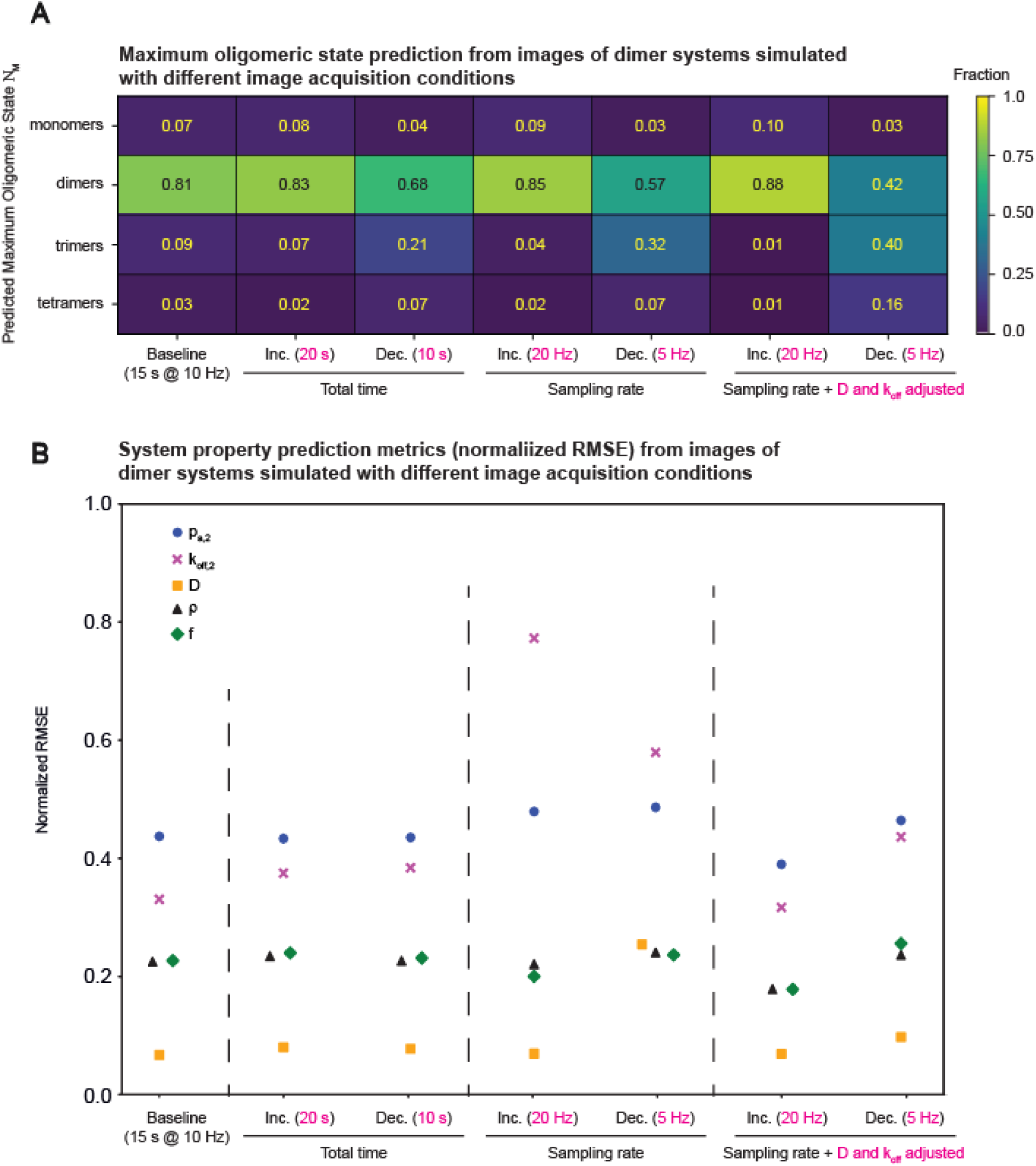
Deep-FISIK is largely robust to variations in imaging conditions as long as they suit the SM system properties. **(A)** Classification accuracy when applying the images classification model (same model as in Fig. 3) to image datasets of dimeric systems generated using various image acquisition conditions: increased or decreased total time, increased or decreased sampling rate, or altered sampling rate combined with compensatory changes in *D* and *k*_off_. All systems are simulated to be dimeric, but the classification model can predict from monomeric to tetrameric. Dataset size = ∼600 graphs per image acquisition condition. **(B)** Normalized RMSE plot for the SM system properties predicted by the images Ν_M_ = 2 regression model (same as in Fig. 3) for image datasets of dimeric systems generated using different image acquisition conditions (same dataset as in A).

The effect of altering the sampling rate was more complex. In terms of the classification task, faster sampling, i.e., more data, improved model performance, while slower sampling, i.e., less data, deteriorated model performance (Fig. 5A). For the regression task, the effects were multi-faceted and much stronger than the effect of changing the total acquisition time. Specifically, for both faster and slower sampling, *k*_off,2_ prediction was heavily negatively impacted. *D* prediction was also heavily negatively impacted for slower sampling. The prediction of *p*_a,2_ was slightly negatively impacted in both scenarios. The properties *ρ* and *f* were largely robust and not affected by changes in sampling rate.

The strong effect of changing the sampling rate on predicting *k*_off,2_ and *D* was to be expected, as the sampling time step (Δt) affects how these properties manifested themselves in the observed system: *D* was represented by the expected (1D) displacement 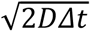 and *k*_off_ was represented by the dissociation probability per time step *k*_off_*Δ*t*. Thus, when the sampling rate was changed, the prediction range of displacements shifted from 0.0316-0.3162 µm (D in the range 0.005–0.5 µm^2^/s (Table 1) and baseline sampling of 10 Hz) to 0.0447-0.4472 µm (5 Hz sampling) or to 0.0224-0.2236 µm (20 Hz sampling). Similarly, the prediction range of dissociation probability per time step shifted from 0.02-0.5 (*k*_off_ in the range 0.2-5/s (Table 1) and baseline sampling of 10 Hz) to 0.04-1.0 (5 Hz sampling) or to 0.01-0.25 (20 Hz sampling). Thus, changing the sampling rate resulted in a situation similar to the out-of-range scenarios investigated above (Fig. S7, S8) for these two properties.

To test whether this was indeed the case, we altered the ranges of *D* and *k*_off_ in the cases of 5 Hz and 20 Hz sampling to return the prediction ranges for 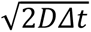 and *k*_off_*Δ*t* to the training range. These changes were in fact biologically/experimentally meaningful, as in an experiment one adapts the sampling rate to the system dynamics/kinetics, where systems with faster dynamics/kinetics would require faster sampling, while slower sampling suffices for systems with slower dynamics/kinetics. For 5 Hz sampling (Δ*t* = 0.2), we shifted the ranges of *D* and *k*_off_ down to 0.0025-0.25 µm^2^/s and 0.1-2.5/s, respectively. For 20 Hz sampling (Δ*t* = 0.05), we shifted the ranges of *D* and *k*_off_ up to 0.01-1 µm^2^/s and 0.4-10/s, respectively. With this, both the classification and prediction accuracies in the 20 Hz scenario returned to their baseline values (Fig. 5A, B). In fact, the classification accuracy with 20 Hz sampling was even better than the baseline accuracy (Fig. 5A), probably because of the increased amount of data with faster sampling. For the 5 Hz sampling with property shifting, the prediction accuracies improved as well, although they remained a little worse than the baseline (Fig. 5B), probably because of the reduced amount of data with slower sampling. At the same time, the classification accuracy deteriorated further in this case, with many systems getting misclassified as trimeric (Fig. 5A). This trend resembles the trend of reducing the total acquisition time, where also many systems were misclassified as trimeric. This suggests that the misclassification in this case stems largely from not capturing slow dissociation events; in other words, slower system kinetics require longer total acquisition time in addition to slower sampling.

Overall, these results indicate that Deep-FISK is generally robust to changes in imaging conditions, as long as the imaging conditions are consistent with the system dynamics and kinetics and the underlying space of property values remains consistent with the training range.

## Discussion

We developed in this work a GNN-based framework, Deep-FISIK, to characterize, from SMI data, the interaction kinetics of molecules (particularly receptors) in their native cellular environment. In the context of homotypic interactions, Deep-FISIK first predicts the maximum oligomeric state of the system, followed by predicting the relevant interaction kinetics and other SM system properties. If the maximum oligomeric state of a system is known a priori, then the first task may be skipped, or this prior knowledge can be used as means to validate Deep-FISIK’s predictions.

A particular strength of Deep-FISIK is that it does not require particle tracking, rather only particle detections in SMI image series. This is important because, as our proof-of-principle testing has shown, which also matches our earlier study (which was nevertheless tracking-based)^46^, the prediction accuracy of all system properties increases as the labeled fraction of molecules increases, at least up to a labeled fraction of ∼0.7-0.8. This is the case even for prediction from images (and not only from pure simulations), even though image data have limited resolution, confounding the detection of molecular interactions. As accurate detection is more compatible with higher labeling densities than accurate tracking, SMI data acquired for Deep-FISIK can be labeled at higher densities (i.e., higher fractions) to increase the accuracy of SM system property prediction.

An important next test for Deep-FISIK would be to apply it to and validate it on experimental data. A complication with experimental data is that the ground truth is generally unknown. Therefore, as discussed above, for this next level of validation it might be necessary to combine Deep-FISIK with various system perturbations that affect molecular interactions in well-defined manners and then verify that Deep-FISIK captures the expected changes. Such studies will establish the versatility and utility of Deep-FISIK for quantifying receptor interaction kinetics and other SM system properties, in order to, ultimately, elucidate the spatiotemporal regulation of receptor and cell signaling.

### Limitations of the study

The Deep-FISIK GNN models trained in this study are limited to one-color SMI data, and thus to one-receptor systems undergoing homotypic interactions. For broader applicability, Deep-FISIK must be generalized to multi-receptor systems, where heterotypic interaction kinetics are predicted. This involves expanding the model employed to generate data for training, including the labeling of different receptors with different “colors”. The graphs as input to Deep-FISIK must be generalized as well. Note that the multi-color generalization may be also applied to a one-receptor system undergoing homotypic interactions, but where the system is probed with multi-color SMI. In addition, the current graph generation process for Deep-FISIK connects detected particles between consecutive frames only. Thus, it cannot handle low SNR cases where molecules may temporarily disappear (due to detection failure) and then be detected again. For this, the graph generation process must be generalized to connect detected particles between non-consecutive frames, thus allowing gap closing to compensate for missed detections^60^. This would extend the range of SNRs to which Deep-FISIK is applicable.

Finally, Deep-FISIK is currently limited to molecules diffusing and interacting in 2D. For broader applicability, it must be extended to 3D. This involves primarily generalizing the model of molecular diffusion and interactions used for data generation and model training, and naturally the graph generation process to take the z-coordinate into account when calculating the distance between detected particles.

## Resource availability

- Deep-FISIK and other custom code written for this study can be found online at https://github.com/khainguyen20/DeepFISIK.
- Example data can be also found online at https://github.com/khainguyen20/DeepFISIK.
- Any additional information required to reanalyze the data reported in this paper is available from the lead contact upon request.

## Acknowledgments

We would like to thank Dr. Jesus Vega-Lugo for help with generating simulations and synthetic movies, Dr. Aparajita Dasgupta for help with running u-track, Dr. Jaime Guerrero for supplying the VEGFR2 experimental diffusion coefficient distribution, and Dr. Satwik Rajaram and the Rajaram lab for feedback on the figures and data presentation. We would also like to thank the UT Southwestern BioHPC high performance computing cluster for computational infrastructure and support. This work was supported by funding from the National Institutes of Health/National Institute of General Medical Sciences (R35 GM119619) (PIs: K. Jaqaman and A. Dasgupta) and from the University of Texas Southwestern Medical Center Endowed Scholars Program to K. Jaqaman.

## Author contributions

K.N., conceptualization, methodology, software, visualization, and manuscript writing. K.J., conceptualization, methodology, resources, funding acquisition, supervision, visualization, and manuscript writing.

## Supplemental information titles and legends

Document S1 contains 8 supplemental figures:

In addition, there are 8 movies, with the following legends:

**Movie S1. SMI of high SNR simulation.** Video of synthetic SMI data, consisting of 150 time points at a sampling rate of 10 Hz (left) with detections overlayed (right). Each displayed area is

9.0 x 9.0 μm^2^. Synthetic data were simulated with the following parameters (parameter of interest underlined): *p*_a,2_ = 0.1, *k*_off,2_ = 2.5 s^−1^, *D* = 0.25 μm^2^/s, *ρ* = 5 # molecules/µm^2^, *f* = 0.5, <u>SNR = 7</u>.

**Movie S2. SMI of low SNR simulation.** Video of synthetic SMI data, consisting of 150 time points at a sampling rate of 10 Hz (left) with detections overlayed (right). Each displayed area is

5.8 x 5.8 μm^2^. Synthetic data were simulated with the following parameters (parameter of interest underlined): *p*_a,2_ = 0.1, *k*_off,2_ = 2.5 s^−1^, *D* = 0.25 μm^2^/s, *ρ* = 5 # molecules/µm^2^, *f* = 0.5, <u>SNR = 4</u>.

**Movie S3. SMI of high-density simulation.** Video of synthetic SMI data, consisting of 150 time points at a sampling rate of 10 Hz (left) with detections overlayed (right). Each displayed area is 5.3 x 5.3 μm^2^. Synthetic data were simulated with the following parameters (parameters of interest underlined): *p*_a,2_ = 0.1, *k*_off,2_ = 2.5 s^−1^, *D* = 0.25 μm^2^/s, *ρ* = 10 # molecules/µm^2^, *f* = 1, SNR = 5.5.

**Movie S4. SMI of low-density simulation.** Video of synthetic SMI data, consisting of 150 time points at a sampling rate of 10 Hz (left) with detections overlayed (right). Each displayed area is 21.1 x 21.1 μm^2^, cropped from 42.8 x 42.8 μm^2^. Synthetic data were simulated with the following parameters (parameters of interest underlined): *p*_a,2_ = 0.1, *k*_off,2_ = 2.5 s^−1^, *D* = 0.25 μm^2^/s, *ρ* = 0.1 # molecules/µm^2^, *f* = 0.1, SNR = 5.5.

**Movie S5. SMI of high affinity simulation.** Video of synthetic SMI data, consisting of 150 time points at a sampling rate of 10 Hz (left) with detections overlayed (right). Each displayed area is 9.3 x 9.3 μm^2^. Synthetic data were simulated with the following parameters (parameters of interest underlined): *p*_a,2_ = 0.2, *k*_off,2_ = 0.2 s^−1^, *D* = 0.25 μm^2^/s, *ρ* = 5 # molecules/µm^2^, *f* = 0.5, SNR = 5.5.

**Movie S6. SMI of low affinity simulation.** Video of synthetic SMI data, consisting of 150 time points at a sampling rate of 10 Hz (left) with detections overlayed (right). Each displayed area is 7.9 x 7.9 μm^2^. Synthetic data were simulated with the following parameters (parameters of interest underlined): *p*_a,2_ = 0.001, *k*_off,2_ = 5 s^−1^, *D* = 0.25 μm^2^/s, *ρ* = 5 # molecules/µm^2^, *f* = 0.5, SNR = 7.

**Movie S7. SMI of fast-moving simulation.** Video of synthetic SMI data, consisting of 150 time points at a sampling rate of 10 Hz (left) with detections overlayed (right). Each displayed area is 8.7 x 8.7 μm^2^. Synthetic data were simulated with the following parameters (parameter of interest underlined): *p*_a,2_ = 0.1, *k*_off,2_ = 2.5 s^−1^, *D* = 0.5 μm^2^/s, *ρ* = 5 # molecules/µm^2^, *f* = 0.5, SNR = 7.

**Movie S8. SMI of slow-moving simulation.** Video of synthetic SMI data, consisting of 150 time points at a sampling rate of 10 Hz (left) with detections overlayed (right). Each displayed area is 6.6 x 6.6 μm^2^. Synthetic data were was simulated with the following parameters (parameter of interest underlined): *p*_a,2_ = 0.1, *k*_off,2_ = 2.5 s^−1^, *D* = 0.005 μm^2^/s, *ρ* = 5 # molecules/µm^2^, *f* = 0.5, SNR = 7.

## Methods

### Key resources table

| REAGENT or RESOURCE | SOURCE | IDENTIFIER |
| --- | --- | --- |
| Software and algorithms |  |  |
| Deep-FISIK | This work | https://github.com/khainguyen20/DeepFISIK |
| Python 3.8.18 | Python | https://www.python.org/ |
| pytorch 2.1.0+cu121 | Pytorch | https://pytorch.org/ |
| torch_geometric 2.4.0 | PyG | https://pytorch-geometric.readthedocs.io/en/latest/ |
| numpy 1.24.4 | Numpy | https://numpy.org/ |
| matplotlib 3.7.2 | Matplotlib | https://matplotlib.org/ |
| scikit-learn 1.3.2 | Scikit-learn | https://scikit-learn.org/stable/ |
| FISIK | de Oliveira<br>et al.<sup>46</sup> | https://github.com/kjaqaman/FISIK |

### Model architecture

The architecture of the Deep-FISIK GNN model is comprised of encoder, message-passing, and decoder blocks (Fig. S1A). The encoder block expands the dimensions of the node and edge features. It is composed of a series of multilayer perceptrons followed by GeLU activation and batch normalization. The dimension after embedding was determined via hyperparameter tuning (Fig. S2). The message-passing block passes information between the target node *i* and source node *j* at the *l*th layer through MHA^51–53^, thus updating the node features (Fig. S1B) (See next sections for details). It is repeated L times, with L determined via hyperparameter tuning (Fig. S2). Lastly, after pooling all of the node features together using percentile pooling^67^ (10^th^, 20^th^…90^th^ percentiles) in order to obtain graph-level predictions, the decoder block predicts either the maximum oligomeric state of the system Ν_M_ (classification task) or the SM system-level properties (regression task; properties: association probability *p*_a,ν_ per oligomeric state ν > 1, dissociation rate constant *k*_off,ν_ per oligomeric state ν > 1, diffusion coefficient distribution mean *D*, receptor density *ρ*, and labeled fraction *f*). Similar to the encoder block, the decoder block is composed of a series of multilayer perceptrons followed by GeLU activation and batch normalization.

### Multi-head attention (MHA)

In MHA^51–53^, the node and edge features (*h* and *e*, respectively) are turned into queries, keys, values, and edge matrices:

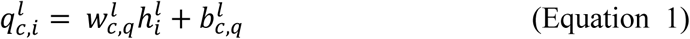

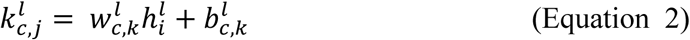

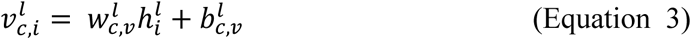

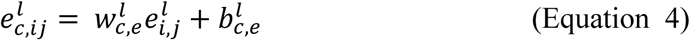

where the *w*’s are weight matrices, the *b*’s are bias terms, *c* is the attention head index and *l* is the message-passing layer index (*l* = 1, 2, …, *L*, where *L* is determined via hyperparameter tuning). The FISIK GNN models use 6 attention heads (c = 1, 2, … C, where C = 6).

With these matrices, the node features per layer are updated as follows:

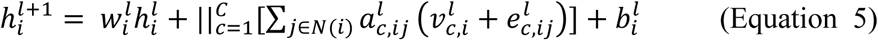

where

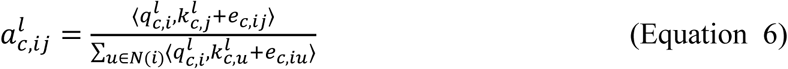

and

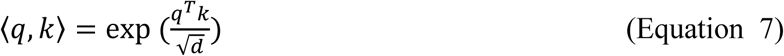

In Eq. 7, *d* is the attention head dimension (e.g., with 96 node features and C = 6 attention heads, *d* = 96/6 = 16). In Eq. 5, the *w*’s and b’s are again weight matrices and bias terms, respectively.

### Intensity-based edge features

To calculate the intensity-based edge features, let *I*_i_ and *I*_j_ be the intensities of particle/node *i* at timepoint *t* and particle/node *j* at timepoint *t*+1. Define Δ*I*_ij_ = *I*_j_ – *I*_i_. Let *I*_mono_ be the average intensity of a monomer (a single fluorophore). If Δ*I*_ij_ ≈ 0, this indicates that the two connected nodes most likely have the same oligomeric state, suggesting that no association or dissociation event has taken place. If Δ*_Iij_* ≈ Δ*ν* × *I_mono_*, Δ*ν* > 0 (i.e., 1, 2, 3, …), this indicates that the oligomeric state of node *j* is Δ*ν* units higher than that of node *i*, suggesting an association event (e.g., for Δ*ν* = 1, node *j* could be a dimer and node *i* a monomer). If, on the other hand, Δ*I_ij_* ≈ Δ*ν* × *I_mono_*, Δ*ν* < 0, this indicates that the oligomeric state of node *j* is *ν* units lower than that of node *i*, suggesting a dissociation event. Mathematically, the intensity-based features – probability of no oligomeric state change *p*_0_, probability of an association (merge) event *p*_m_, and probability of a dissociation (split) event *p*_s_ – are defined as follows (see Fig. 1A inset for visual representation):

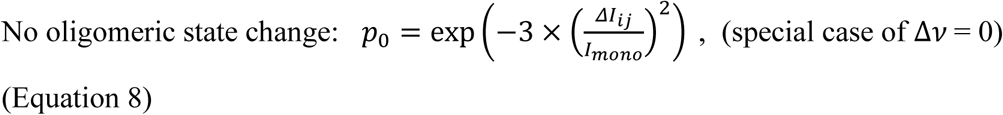

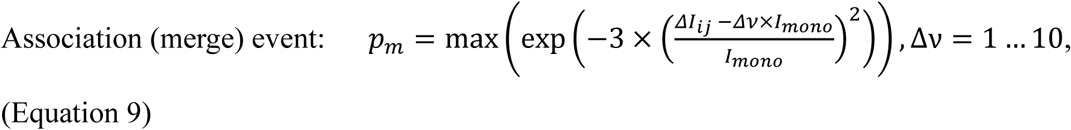

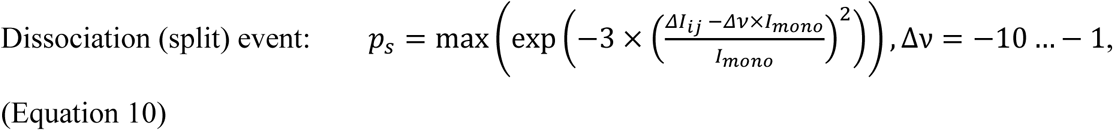

Eqs. 9 and 10 allow for oligomeric state differences Δ*_ν_* up to 10. This is sufficient for molecular systems that form oligomeric states up to 11-mer (monomer + decamer → 11-mer). If a system is known to form higher-order oligomers, then Eqs. 9 and 10 can be expanded accordingly.

### Simulation of synthetic SMI data of 2D molecular movement and interactions

To generate training and validation datasets for Deep-FISIK, we employed a stochastic model of molecular diffusion and interactions in 2D, part of which was also substoichiometric labeling to mimic single-molecule sampling^46^ (Fig. 2A). For pure simulation datasets and models, molecule positions were taken directly from the simulations and used to generate the GNN model input graphs. For image datasets and models, the simulated molecule positions per timepoint were the starting point to generate image series^59^, in which particles were subsequently detected, and then the GNN model input graphs were generated from the detections. This section describes the simulations and image generation, while the next section describes the detection process.

In the simulations, molecules (all of one type) with density *ρ* undergo free diffusion, where each molecule has a diffusion coefficient *D*_SM_ drawn from a normal distribution with specified mean *D* and standard deviation *σ*_D_ = 0.15*D* . When a monomer and an oligomer of oligomeric state ν (ν = 1, 2, …; note that an oligomer with state ν = 1 is a monomer) encounter each other (i.e., they are < 10 nm apart), they have a probability *p*_a,ν+1_ to associate with each other and form an oligomer of state ν +1. The maximum oligomeric state allowed in a system is defined as N_M_ (in other words, *p*_a,ν_ = 0 for ν > Ν_M_). A monomer can dissociate from an oligomer of state ν ≥ 2 with dissociation rate constant *k*_off,ν_, thus forming a monomer and an oligomer of size ν-1. To mimic substoichiometric labeling, a fraction *f* of the molecules is labeled, leaving the remaining fraction (1- *f*) invisible. The intensity of an individual fluorophore *I*_fl_ ∼ *N*(*μ*_fl_, *σ*_fl_), where *μ*_fl_ = 1 and *σ*_fl_ = 0.15, representing the intensity fluctuations of an individual fluorophore over time. The simulations were performed in an area of 30×30 µm^2^ for 15 s (after an initialization period of 10 s) using a simulation time step of 0.01 s, and were subsequently subsampled to 10 Hz (i.e., sampling step Δ*t* of 0.1 s) to match experimental SMI data^59^.

In our training and validation datasets, *D*, *ρ*, *f*, Ν_M_, *p*_a,ν_, and *k*_off,ν_ (ν=2… Ν_M_) were generated such that each simulation has a different combination of these SM system properties (model parameters). Note that *p*_a,ν_ and *k*_off,ν_ were different for the different ν within a simulation. The ranges for these properties, motivated by prior studies^9,59,65,66^, are shown in Table 1 and Fig. 2A.

To decouple the number of molecules in a simulation from *ρ* and to speed up computation, we did not use the full simulated area for generating graphs representing each simulation. Rather, we utilized a sub-area from the middle of each simulated area, so that the number of molecules per timepoint in the graph was between 50 and 250 molecules, regardless of the density *ρ*.

From these pure simulations, synthetic images were then generated to mimic experimental SMI data^59^. This was achieved by placing a point spread function (PSF), approximated by a Gaussian with standard deviation σ_PSF_, at the location of each labeled molecule, and then adding background offset and noise. The image boundaries were also padded with empty areas (10σ_PSF_ on all sides) to avoid molecules falling on the boundary and becoming undetectable. The synthetic image generation parameters are shown in Table 2.

**Table 2.** Synthetic Image Generation Parameters.

| Parameter | Value/value range |
| --- | --- |
| Signal-to-noise | 4-7 |
| Pixel size | 81 nm x 81 nm |
| $\sigma_{\text{PSF}}$ | 122 nm |

### Particle detection in SM image series

Particle detection in the generated images was performed using u-track^60^, using the iterative Gaussian-Mixture Model algorithm, with the parameters shown in Table 3.

**Table 3.** Particle Detection Parameters.

| Parameter | Value |
| --- | --- |
| Rolling window | 1, 3, 5 |
| $\alpha_{\text{local,max}}$ | 0.1 for rolling window 1<br>0.05 for rolling window 3<br>0.05 for rolling window 5 |
| $\alpha_{\text{residuals}}$ | 0.1 |
| $\alpha_{\text{amplitude}}$ | 0.1 |
| $\alpha_{\text{distance}}$ | 0.1 |

The α-values were chosen based on visual assessment of the detection results with the goal of minimizing both false positives (superfluous detections) and false negatives (missed particles).

### Graph generation

Starting with the particle positions per timepoint – whether directly from the simulations (pure simulations dataset) or upon detection in the synthetic images (images dataset) – the input graph to Deep-FISIK was generated by taking each particle as a node and connecting the nodes in different timepoints to make graph edges (Fig. 2B). The generated graphs were directed, with information flowing from the first time point to the last time point. Particles in different timepoints were connected in the graph if the distance between them was smaller than a user-defined search radius. In our study, the search radius was taken as 1.5 µm, reflecting the maximum displacement of a molecule given the range of diffusion coefficients employed in the simulation and the simulation time step (the extreme of the displacement distribution given the largest possible D_SM_). In the current implementation, connections between particles were limited to consecutive timepoints, i.e., there was no gap closing to account for temporary particle disappearance^60^. Graph building can however be generalized to connect particles between non-consecutive timepoints if needed. To speed up computation, each graph was trimmed by retaining only the 3 shortest edges (3 closest distances between nodes) for each source node and for each target node and taking the union of these edges.

For both model training and validation, the node features (x- and y-coordinates and intensity) were min-max normalized and the distance edge feature was normalized by the search radius. The intensity-related edge features were by definition normalized between 0 and 1 (Eqs. 1-3). Each graph was labeled with its system-level properties: 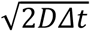, *ρ*, *f*, Ν_M_, *p*_a,ν_, and *k*_off,ν_*Δ*t* (ν = 2… Ν_M_). Using the expected (1D) displacement 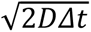 instead of *D*, and similarly using the dissociation probability per time step *k*_off_*Δ*t* instead of *k*_off_, accounted for the role that the sampling time step Δ*t* played in how these system properties manifested themselves in the observed system.

### Model training

For the classification models (pure simulations and images), we used 20,000 graphs (5,000 per oligomeric state). For the Ν_M_ = 2 regression models (pure simulations and images) and Ν_M_ = 4 regression model (pure simulations), we used 30,000 graphs. For the Ν_M_ = 3 model (pure simulations), we used first 30,000 graphs and then 60,000 graphs. For each model training, we split the dataset into 80% for training and 20% for testing. The batch size was 100 graphs for the Ν_M_ = 2 model (pure simulations and images) and 200 graphs for all other models. We used Adam optimizer, with a learning rate of 10^−4^ that was decreased by a factor of 0.4 every 15 epochs. We trained each model for a maximum of 100 epochs and did hyperparameter tuning with various hyperparameters (Fig. S2).

The loss functions were cross-entropy loss (ℒ*_CE_*) and sum of mean-squared errors (ℒ*_MSE_*) for the classification task and regression task, respectively, defined as follows:

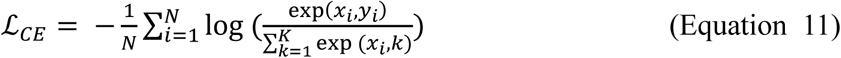

where *i* is the graph index, N is the number of graphs in batch (200), K is the number of classes (4 in our training), *x*_i_ is the ground truth class of graph *i*, and *y*_i_ is the predicted class of graph *i*.

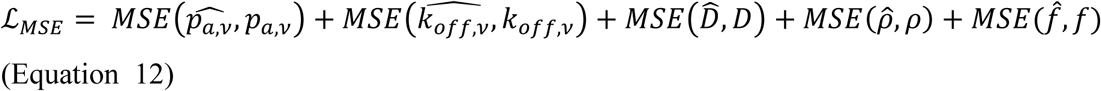

wheri

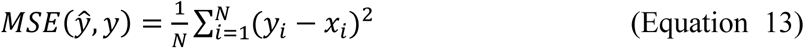

where *i* is the graph index, N is the number of graphs in batch (200), *x*_i_ is the ground truth property value of graph *i*, and *y*_i_ is the predicted property value of graph *i*.

As can be seen from Eq. 12, all properties contribute together to the regression loss function. Therefore, to put all of them on equal footing in the loss function (Eq. 12), we scaled them so that they all had the same maximum value. We tested maximum values of 1, 10, 100, and 1000 and found that a maximum value of 10 yielded the best results.

All models were trained on the UT Southwestern BioHPC Nucleus cluster using a compute node configured with 512 GB RAM and 2x Intel Xeon Gold 6354 processors (36 cores @ 3.0 GHz).

---

**Supplementary Figure S1.**
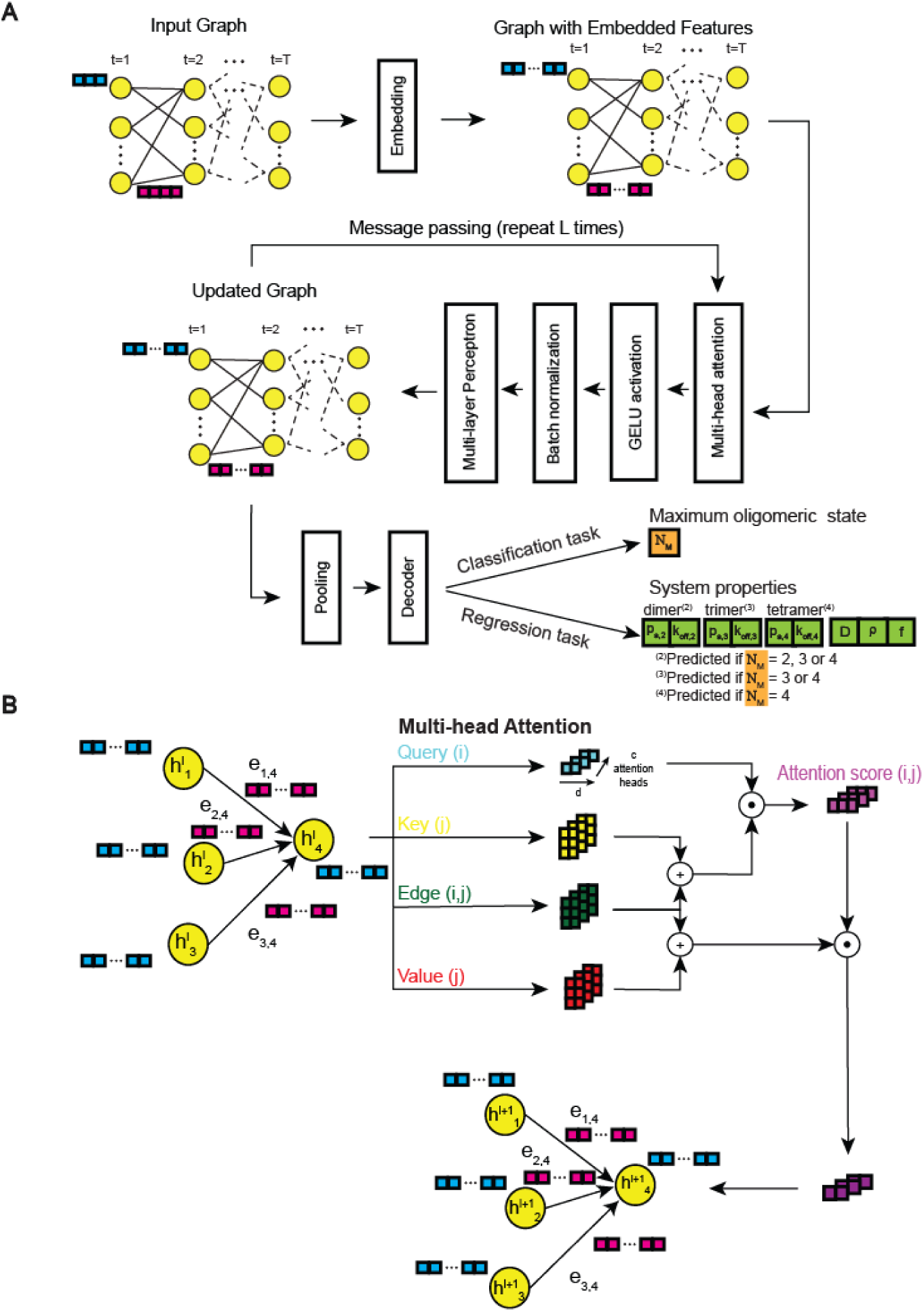
Architecture of the Deep-FISIK models. **(A)** The main layers of the Deep-FISIK GNN models. The node and edge features from the input graph are embedded into a higher dimension that allows for more complex learning. Then, the graph is passed through the message-passing layers L times to update the graph node features based on their spatiotemporal connectivity. Finally, the node features are pooled together using percentile pooling and passed through a decoder to perform either the classification task (determine the maximum oligomeric state Ν_M_ of the system) or the regression task (predict the SM system properties *D*, *ρ*, *f*, *p*_a,2-4_, *k*_off,2-4_). **(B)** Illustration of multi-head attention (MHA), used for message-passing. The node features are turned into queries, keys, and values, which are combined with the edge features and used to calculate attention scores between connected nodes, thus updating the node features. In the schematic, + indicates addition and “dot” indicates dot product.

**Supplementary Figure S2.**
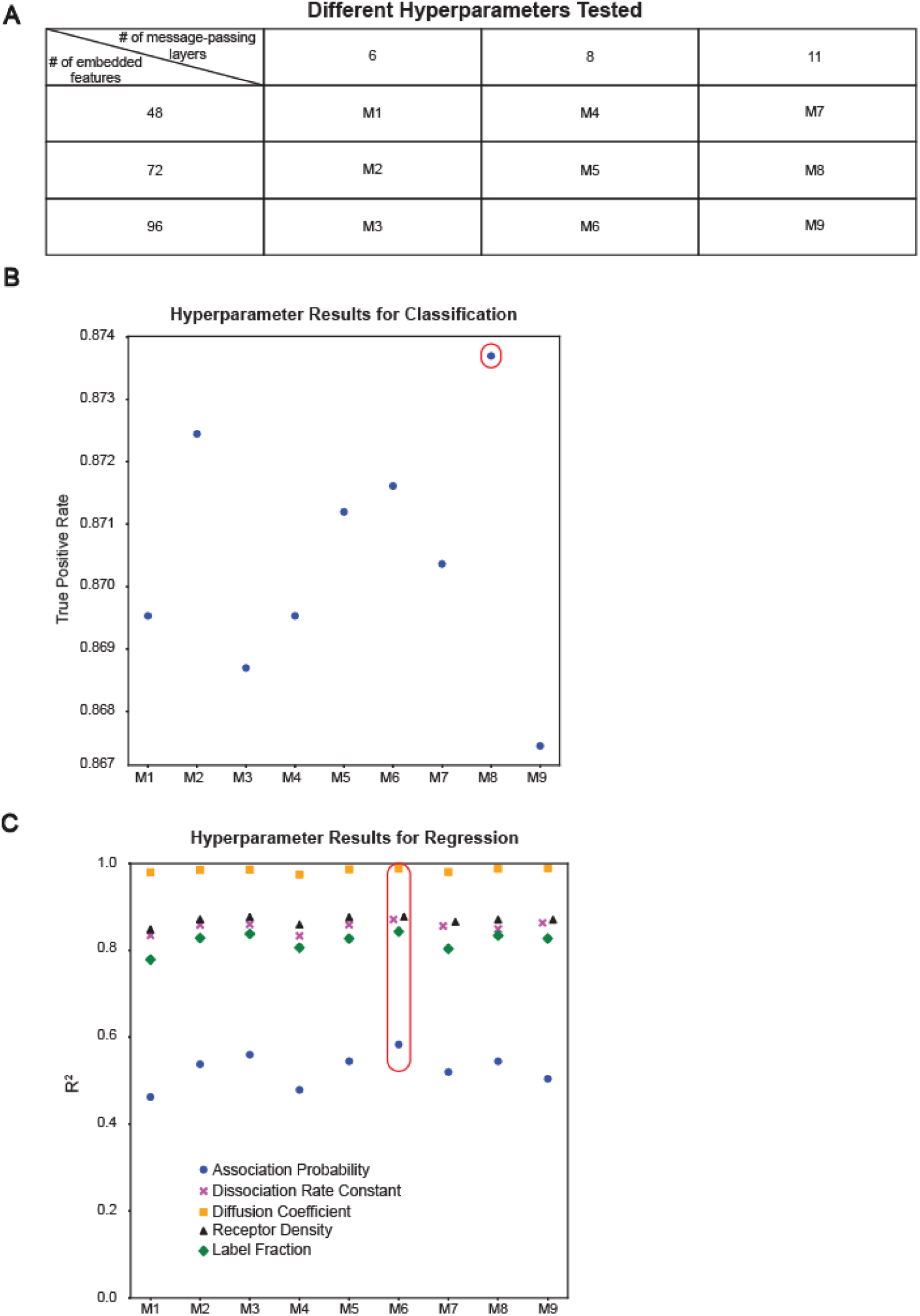
Choosing the best-performing models via hyperparameter tuning. **(A)** Table of hyperparameter combinations used to optimize the Deep-FISIK GNN models for the classification and regression tasks. In total 9 models (M1-M9) with the indicated combinations of number of message-passing layers L and number of embedded features were compared. **(B)** True positive rate in the classification task (20,000 graphs for training and 2,400 for evaluation) using the different GNN model hyperparameters (M1-M9). **(C)** SM system property prediction accuracy (R^2^) in the Ν_M_ = 2 regression task (30,000 graphs for training and 600 for evaluation) using the different GNN model hyperparameters (M1-M9). In B, C, the best performing models are circled in red.

**Supplementary Figure S3.**
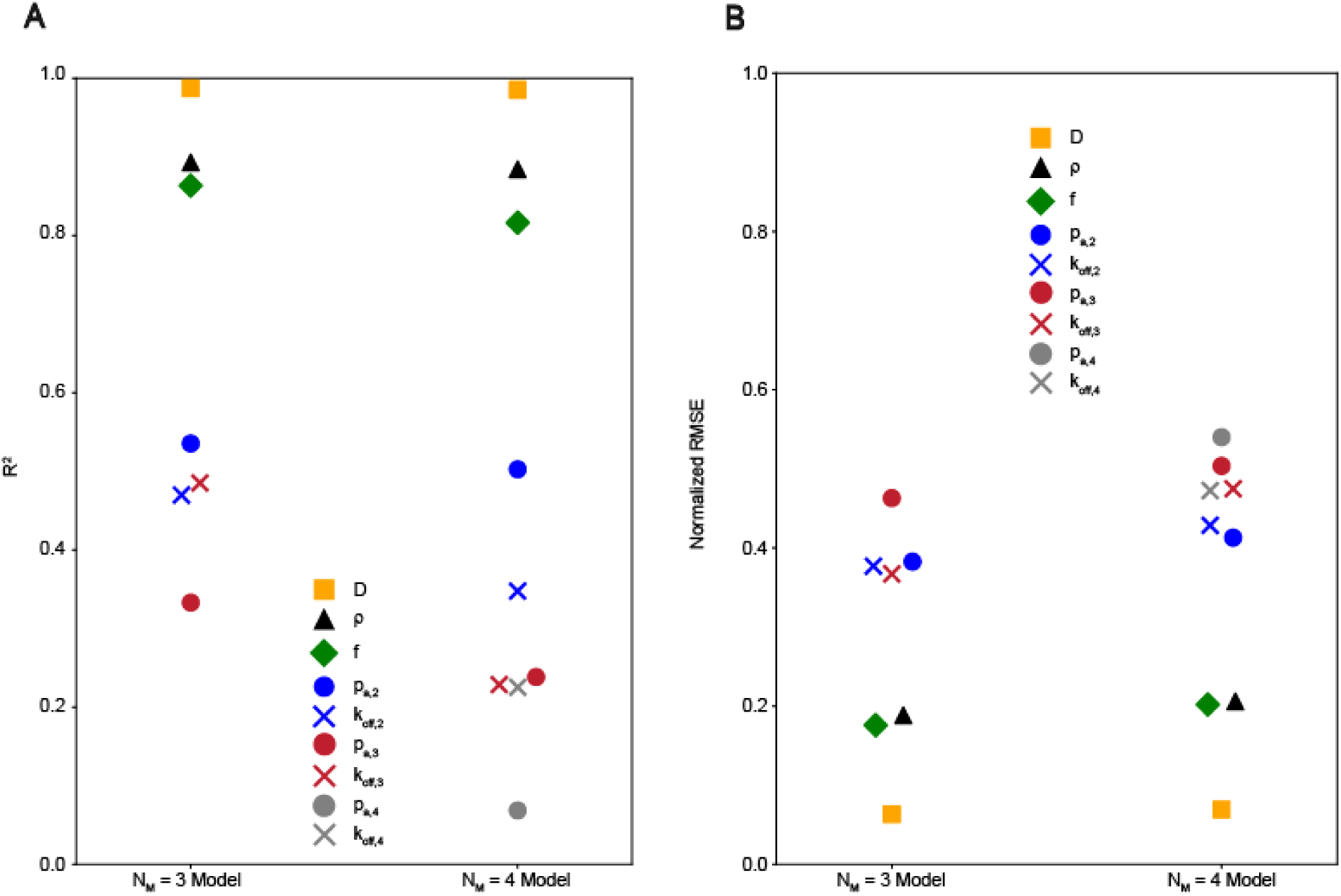
Performance metrics for the pure simulations ΝM = 3 and ΝM = 4 regression models as trained on 30,000 graphs each. **(A)** R ^2^ plot and **(B)** normalized RMSE plot for the various SM system properties for the two models. Each model was evaluated on 600 unseen graphs.

**Supplementary Figure S4.**
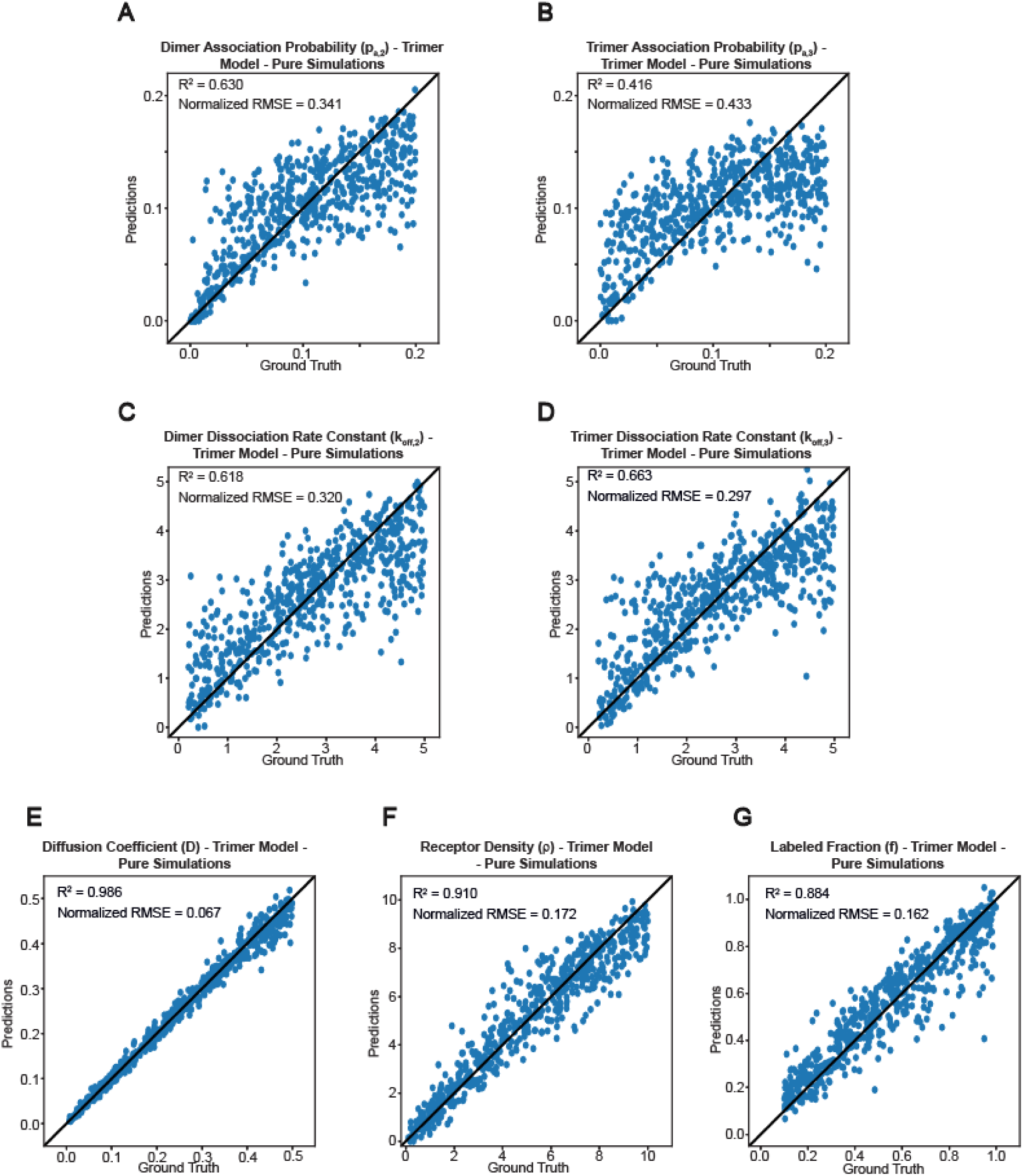
For pure simulated trimeric systems, Deep-FISIK predicts SM system properties with relative error of 7-44% when trained on 60,000 graphs. (A-G) Scatterplots of predicted SM system properties vs. their ground truth values for the pure simulations Ν_M_ = 3 regression model, as obtained by applying the model to unseen trimeric system data. The seven properties are dimer association probability *p*_a,2_ **(A)**, trimer association probability *p*_a,3_ **(B)**, dimer dissociation rate constant *k*_off,2_ **(C)**, trimer dissociation rate constant *k*_off,3_ **(D)**, diffusion coefficient distribution mean *D* **(E)**, molecule density *ρ* **(F)**, and labeled fraction *f* **(G)**. Black straight line in each panel shows the unity line of perfect predictions. The model was trained on ∼60,000 graphs and evaluated on 600 graphs.

**Supplementary Figure S5.**
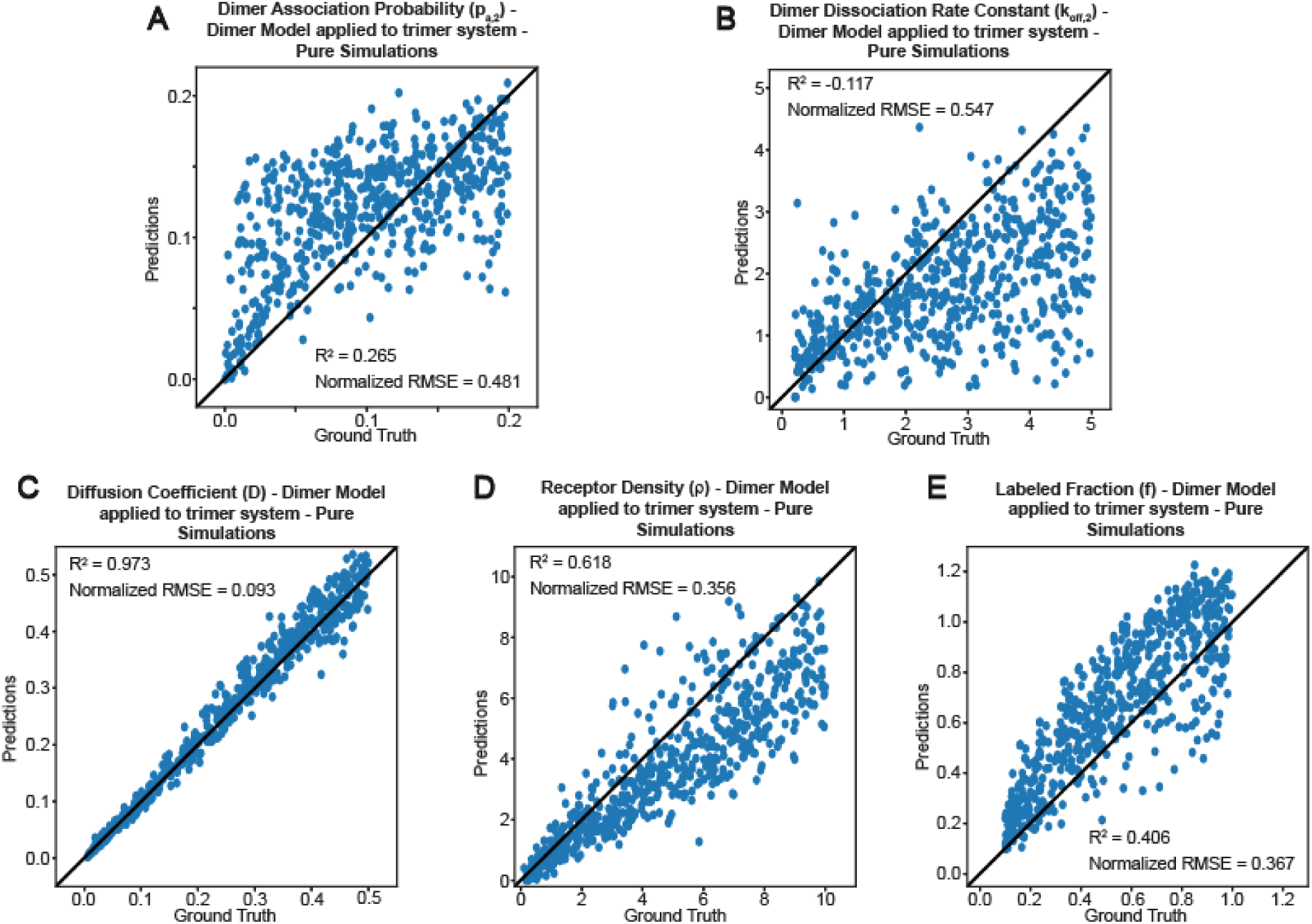
Applying the Ν_M_ = 2 regression model to a trimeric system (emulating a case of underestimating Ν_M_) leads to large errors in most SM system property predictions. (A-E) Scatterplots of predicted SM system properties vs. their ground truth values as obtained by applying the pure simulations Ν_M_ = 2 regression model (same model as in Fig. 2) to unseen data of trimeric systems. The purpose was to determine the consequences of misclassifying a trimeric system as dimeric (i.e., underestimating Ν_M_). The five properties are dimer association probability *p*_a,2_ **(A)**, dimer dissociation rate constant *k*_off,2_ **(B)**, diffusion coefficient distribution mean *D* **(C)**, molecule density *ρ* **(D)**, and labeled fraction *f* **(E)**. Naturally in this case the trimer association probability *p*_a,3_ and trimer dissociation rate constant *k*_off,3_ cannot be predicted. Black straight line in each panel shows the unity line of perfect predictions. The model was tested on 600 graphs.

**Supplementary Figure S6.**
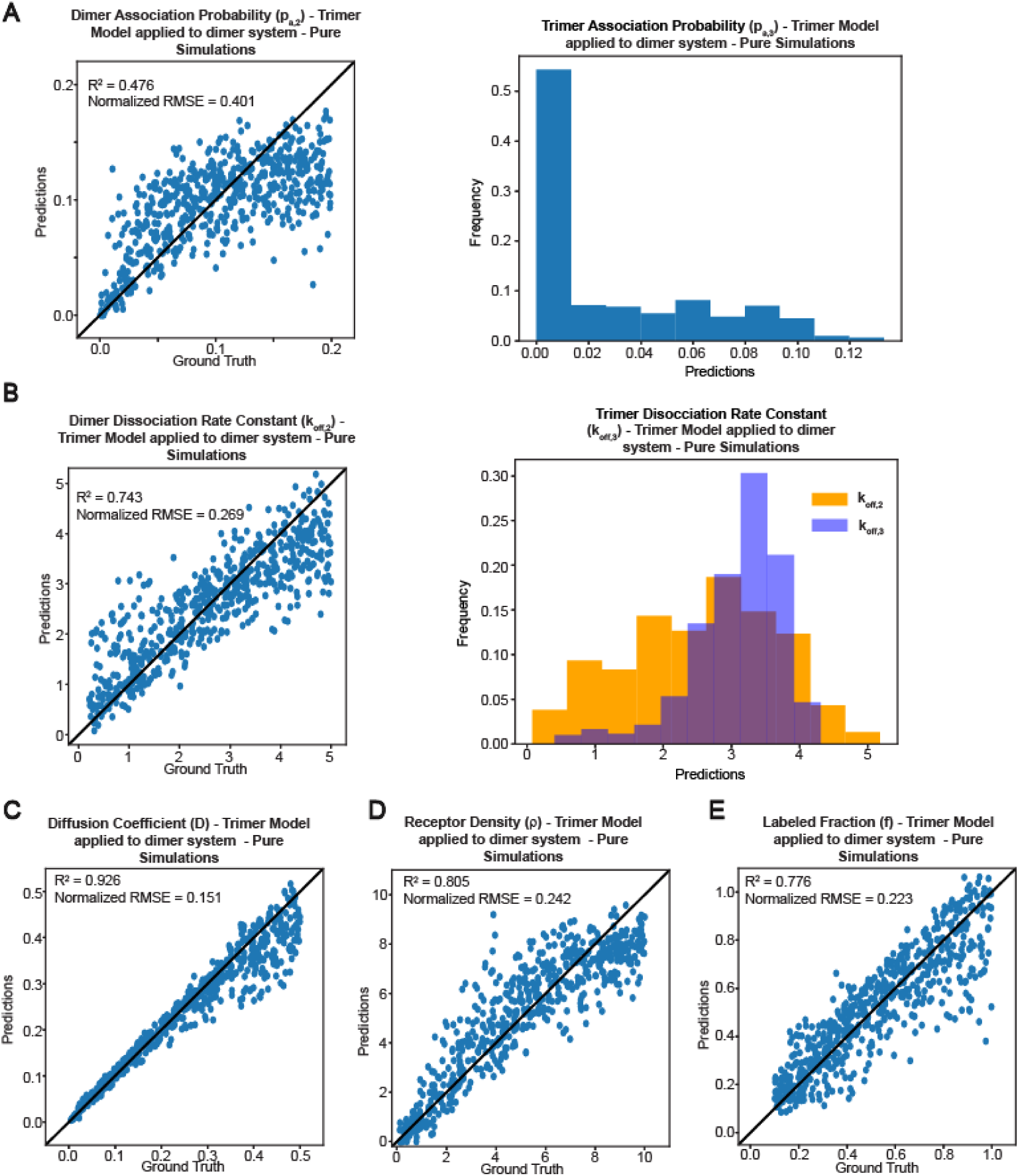
Applying the Ν_M_ = 3 regression model to a dimeric system (emulating a case of overestimating Ν_M_) leads to SM system property predictions slightly worse than (but overall comparable to) those from applying the correct Ν_M_ = 2 regression model. The purpose was to determine the consequences of misclassifying a dimeric system as trimeric (i.e., overestimating Ν_M_). **(A)** Scatterplot of predicted dimer association probability *p*_a,2_ vs. ground truth (left) and distribution of predicted artifactual trimer association probability *p*_a,3_ (right) as obtained by applying the pure simulations Ν_M_ = 3 regression model (same model as in Fig. S4) to unseen data of dimeric systems. **(B)** Same as A but for the predicted dimer (left) and artifactual trimer (right) dissociation rate constants, *k*_off,2_ and *k*_off,3_. In the right panel, the predicted *k*_off,2_ is plotted alongside the artifactual *k*_off,3_ for reference. **(C-E)** Scatterplots of predicted diffusion coefficient distribution mean *D* **(C)**, receptor density *ρ* **(D)**, and labeled fraction *f* **(E)** vs. their ground truth values. Where relevant, black straight line shows the unity line of perfect predictions. The model was tested on 600 graphs.

**Supplementary Figure S7.**
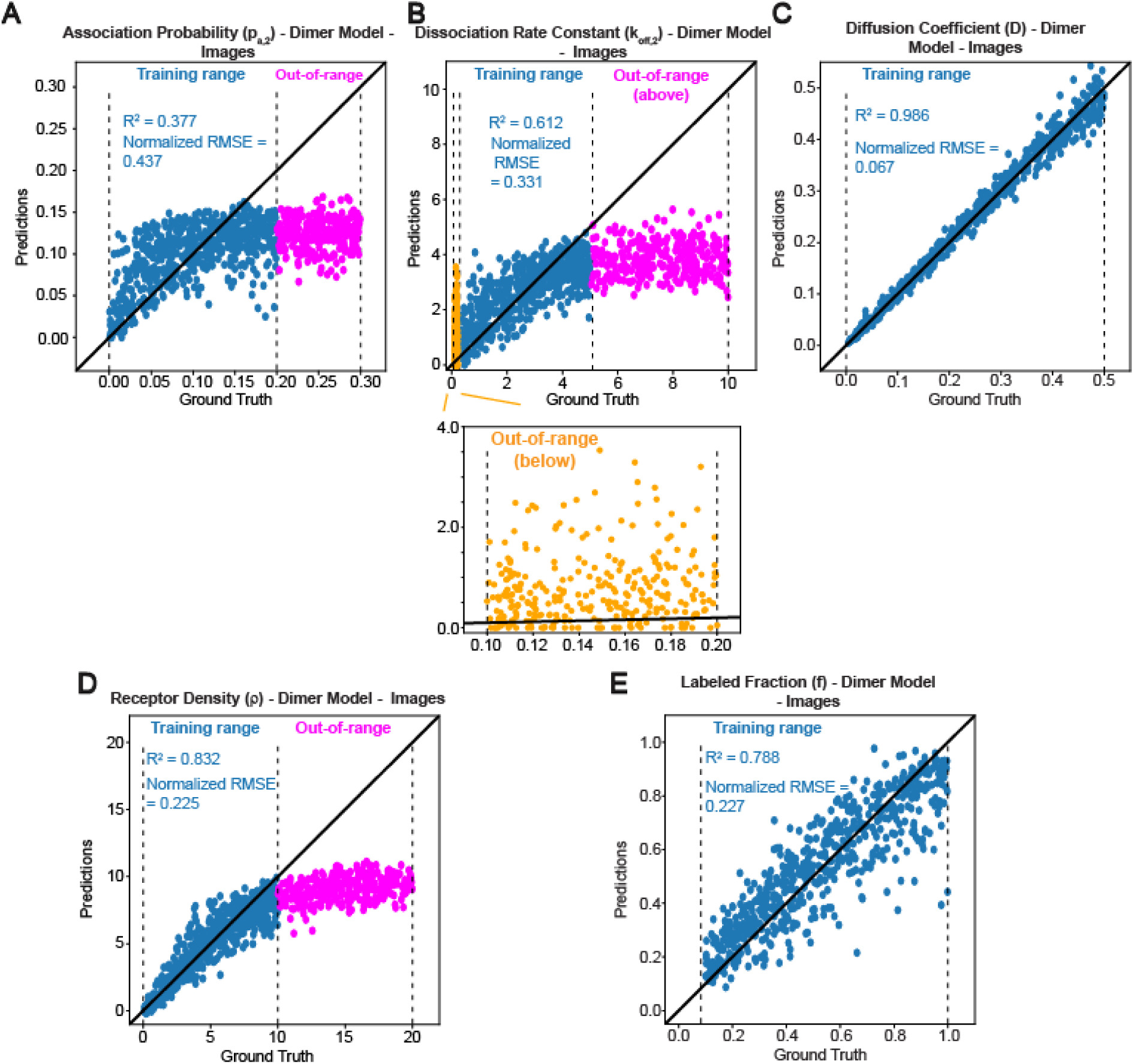
Scatterplots of predicted SM system properties vs. their ground truth values for the images Ν_M_ = 2 regression model. (A-E) Scatter plots were obtained by applying the model (same model as in Fig. 3) to unseen image data of dimeric systems, both within the training range (in-range; blue) and out of range (above-range and below-range; magenta and yellow, respectively). The five properties are dimer association probability *p*_a,2_ **(A)**, dimer dissociation rate constant *k*_off,2_ **(B)**, diffusion coefficient distribution mean *D* **(C)**, molecule density *ρ* **(D)**, and labeled fraction *f* **(E)**. Black straight line in each panel shows unity line of perfect predictions. The in-range data are the same used to generate the diagnostic metrics shown in Fig. 3C-F. The model was evaluated on ∼600 in-range graphs (same as in Fig. 3C-F) and 300 above-range or below-range graphs.

**Supplementary Figure S8.**
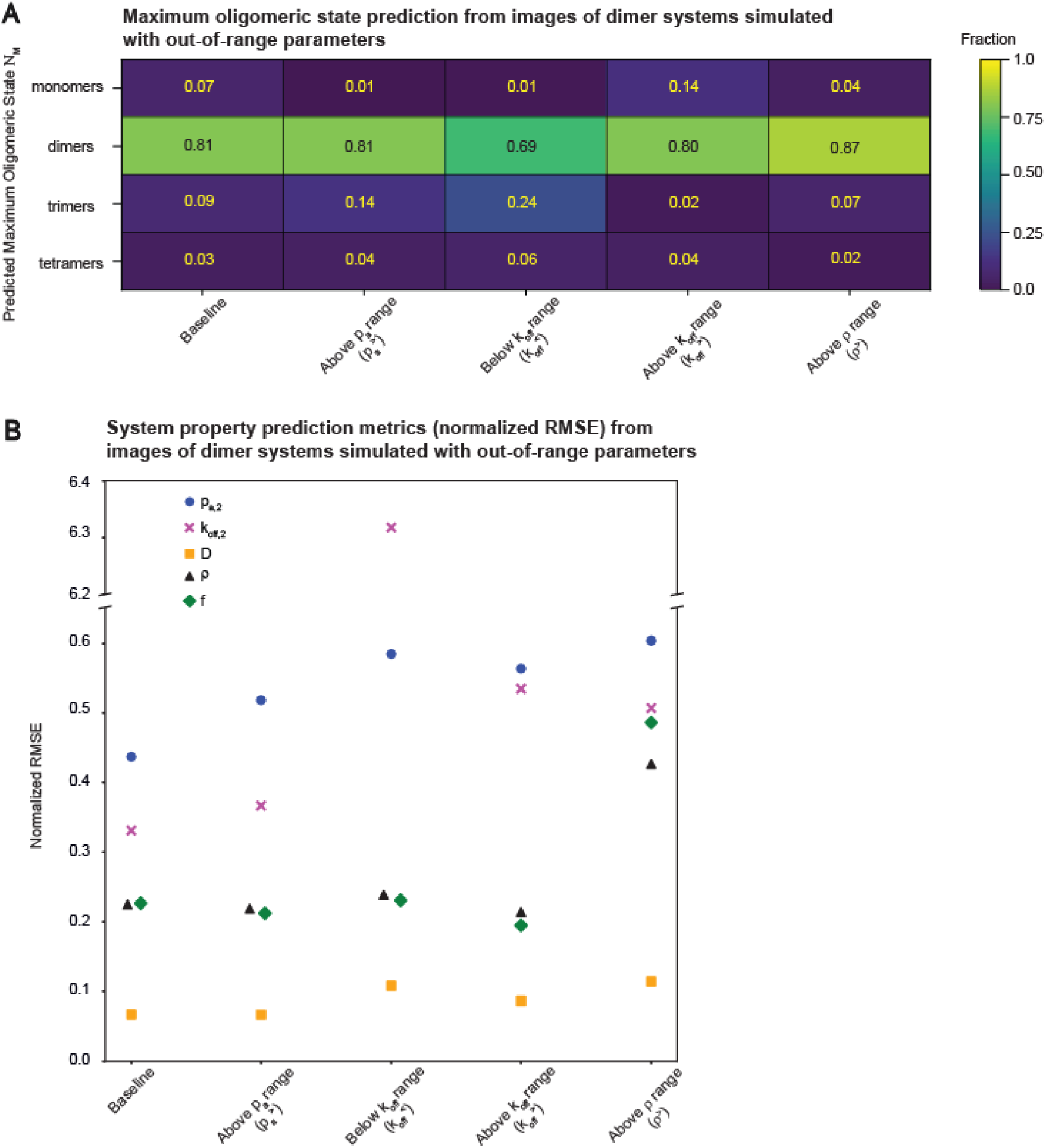
The classification task of Deep-FISIK is largely robust when applied to systems with out-of-range SM system property values, but the regression task cannot extrapolate. **(A)** Classification accuracy when applying the images Ν_M_ = 2 classification model (same model as in Fig. 3) to image datasets of dimeric systems generated using SM system property values outside of the training range (same out of range data shown in Fig. S7): association probability above the training range (*p*_a_^>^), dissociation rate constant below or above the training range (*k*_off_^<^, *k*_off_^>^), and molecule density above the training range (*ρ*^>^). In each case, the other SM system properties are in-range. All simulated systems are dimeric, but the classification model can predict from monomeric to tetrameric. Dataset size = 300 graphs per out-of-range dataset. **(B)** Normalized RMSE plot for the SM system properties predicted by the images ΝM = 2 regression model (same as in Fig. 3) for dimeric system image datasets generated using SM system property values outside of the training range (same dataset as in A).

